# Memory transfer unfolds through rapid shifts in memory stability states during sleep in humans

**DOI:** 10.64898/2026.03.01.708409

**Authors:** Xuemei Li, Makoto Uji, Ryosuke Katsumata, Chisato Suzuki, Kenichi Ueno, Masaki Fukunaga, Sayaka Aritake, R. Allen Waggoner, Masako Tamaki

**Affiliations:** RIKEN Center for Brain Science, Saitama, Japan; Chiba University, Chiba, Japan; National Institute for Physiological Sciences, Okazaki, Japan; Saitama Prefectural University, Saitama, Japan; RIKEN Pioneering Research Institute, Saitama, Japan

**Author notes:** Correspondence to: Masako Tamaki. Contributed equally to this work (Xuemei Li, Makoto Uji).

## Abstract

Sleep benefits learning and memory. Fundamental questions remain regarding whether and how sleep transfers memories for adaptive behavior in humans. We demonstrate that declarative to procedural memory transfer occurs through rapid network reorganization and memory stability shifts in human participants. Following local processing in motor circuit during slow wave–spindle coupling in nonrapid eye movement (NREM) sleep, multiregional communication during phasic rapid-eye movement (REM) sleep enables transfer. By leveraging a newly developed time-resolved simultaneous ultrahigh-field magnetic resonance spectroscopy and polysomnography, we further reveal that memory state becomes instantaneously unstable during slow wave–spindle coupling then enters a hyperstable state during phasic–REM sleep. Thus, sleep bridges memory systems, utilizing increased instability in NREM sleep, and transferring memory through hyperstabilization in REM sleep for knowledge integration.

## Main Text

Our behaviors are influenced by sequences and regularities. Take grocery shopping. You know exactly where to find vegetables, rice, and then noodles in a shop you regularly visit. Even in a new shop, this task is surprisingly easy if the overall layout of item categories is similar. Yet you may find yourself completely lost when the structure or a rule is altered, unable to locate even a bottle of seasoning. This could also apply to a wide range of activities and tasks, including those requiring fine motor skills in which precise sequences of actions are essential, such as the execution of experimental protocols. A crucial aspect of memory is the ability to extract and store higher-order abstract rules from regularities in sequences. Remarkably, when such a higher-order regularity is shared, information can even be transported across the declarative and procedural memory systems, classically considered distinct (*1*, *2*), to facilitate behaviors, a process referred to as memory transfer (*3*, *4*). Such a dynamic form of memory transformation lies at the core of adaptive cognitive function of the human brain.

A growing body of evidence indicates that sleep is essential for memory transformation (*5*, *6*) and integration across memories (*7–9*). However, it has remained unclear whether, and how, sleep promotes the transfer across memory systems through the extraction of regularities for adaptive behavior in the human brain. At least four controversies remain, leaving the fundamental question of how sleep contributes to memory transfer unresolved. First, animal studies have shown that multi-region communications, including the hippocampus–cortex interactions, considered essential for memory transformation (*10*, *11*), occur specifically during slow-wave sleep of nonrapid eye movement (NREM) sleep (*12*, *13*). However, in humans, connectivity is significantly broken down during slow-wave sleep, and modularity in large-scale brain networks instead emerges (*14*, *15*). Thus, it is unclear how slow-wave sleep alone could process information in a manner sufficient to give rise to memory transfer. Second, it has been suggested that sleep microstructures, such as slow-wave–spindle coupling (*16*) and the phasic and tonic periods of rapid-eye movement (REM) sleep (*17*), are involved in memory processing. However, it remains unknown how these brief neural events or periods could produce network-level modifications required for memory transfer. Third, unstable memory is proposed as a crucial state for memory transfer during wakefulness (*18*). Memory stability is measured behaviorally as reduced susceptibility to interference or disruption in performance on a task by subsequent learning or training (*19*). Interference is considered to emerge due to an unstable memory state that is vulnerable to modification made by new information (*20*). However, how memory stability changes rapidly offline in association with sleep microstructures, when overt behavioral responses are largely absent, has never been investigated. Critically, the fourth controversy concerns the prevailing assumption in major models that NREM sleep is the sole sleep stage supporting memory processing, whether through synaptic upscaling or downscaling (*21*, *22*). The active systems consolidation model beautifully explains how neural mechanisms of slow-wave sleep could support transferring memories from the hippocampus to long-term cortical representations (*5*). Accumulating evidence, however, indicates otherwise that REM sleep plays a critical role in memory, particularly in humans who exhibit well-consolidated NREM–REM sleep cycles (*23*, *24*). One study demonstrated that NREM sleep promotes performance improvement, whereas REM sleep reduces interference in visual learning (*23*). Additional studies have shown that REM sleep enhances hippocampus–neocortical communication and memory (*25–27*). While extensive studies have elegantly demonstrated the effects of cueing during NREM sleep for memory transformation (*28*, *29*), these findings do not provide evidence that the physiological mechanisms during REM sleep were unchanged, and whether NREM and REM sleep collectively take part in memory transfer is unclear.

The present study investigated whether sleep facilitates memory transfer by measuring functional connectivity and memory stability changes in association with sleep microstructures for the first time in human participants to resolve all the above-mentioned controversies. While at present there is no direct way to measure the states of memory stability, previous studies have reported that the excitation-to-inhibition (E/I) ratio, theoretically known as the E/I balance (*30*, *31*), measured by magnetic resonance spectroscopy (MRS), is correlated with memory stability (*32*, *33*). Using MRS, the concentrations of the key excitatory neurotransmitter glutamate and the key inhibitory neurotransmitter gamma-aminobutyric acid (GABA) can be estimated (*34*). One can derive the E/I ratio from the ratio of GABA to glutamate concentrations. High E/I ratios are associated with unstable and fragile memory, and low E/I ratios are associated with robust and stable memory (*32*). Thus, the MRS-based E/I ratio is used as a readout for memory stability. However, a major challenge in MRS is its extremely poor temporal resolution (*35*), which hinders investigations into how memory stability shifts in relation to sleep microstructures. The temporal resolution has been limited to 2–10 min with 3-Tesla MRI (*19*, *36*). In contrast, sleep microstructures, including slow waves, their coupling with sleep spindles, and rapid eye movements, can transiently emerge and then subside within a few seconds or less (*37*, *38*). If these actively drive memory transfer, averaging MRS signals every few minutes could falsely negate any effects specific to these states. Thus, leveraging ultrahigh-field 7-Tesla magnetic resonance imaging (MRI), we developed a time-resolved simultaneous MRS and polysomnography (PSG) method, which maximized the temporal resolution in measuring the concentrations of neurotransmitters. We succeeded for the first time in estimating time-varying changes in the E/I ratios on the scale of seconds during sleep in human participants. This allowed us to identify sleep-state-dependent memory state dynamics that had previously been entirely undetectable.

We demonstrate that memory transfer occurs through distinctive processing during NREM and REM sleep through dynamic network and memory state shifts. In experiment 1, we show that local processing in the motor circuit occurs during NREM sleep. Conversely, during subsequent REM sleep, active transfer occurs, where hippocampus – cortical connectivity increases, resulting in the breaking of declarative and procedural system boundaries. In experiment 2, we demonstrate that the memory state is highly unstable, evidenced by increased E/I ratios in association with slow wave–spindle coupling during NREM sleep. Crucially, the memory state becomes hyperstable, indicated by significantly decreased E/I ratios during subsequent REM-phasic period. Besides these rapid shifts, memory stability gradually increases in correlation with delta power. Collectively, the multi-phase processing that the present study proposes not only resolves the long-standing controversies regarding the role of sleep in memory but also provides a novel general framework for how knowledge can be integrated and generated during sleep for adaptive cognition in humans.

## Results

### Sleep, not wakefulness, promotes declarative-to-procedural memory transfer, using high-order information

In experiment 1, we examined whether sleep facilitates declarative to nondeclarative memory transfer and whether it involves functional connectivity changes associated with sleep microstructures. We trained participants using a two-task paradigm before a 90-min interval with or without sleep (Fig. 1a; see the Two-task paradigm in the Materials and Methods section). The tasks included the finger tapping motor sequence task (hereafter, motor task; see <u>Motor task</u> in the Materials and Methods section), which is a nondeclarative procedural task (*39*, *40*), and the category memory task (hereafter, category task), which is a declarative task. In the motor task, subjects pressed four numeric keys on a standard computer keyboard using their nondominant hand, repeating a twelve-element sequence as quickly and as accurately as possible. In the category task, participants memorized the order of categories of twelve visual stimuli, which were composed of four categories (object, scene, face, and animal). Two tests, the probe and the recall tests, were conducted within the category task to facilitate memorization of the sequence (see <u>Category task</u> in the Materials and Methods section).

**Fig. 1:**
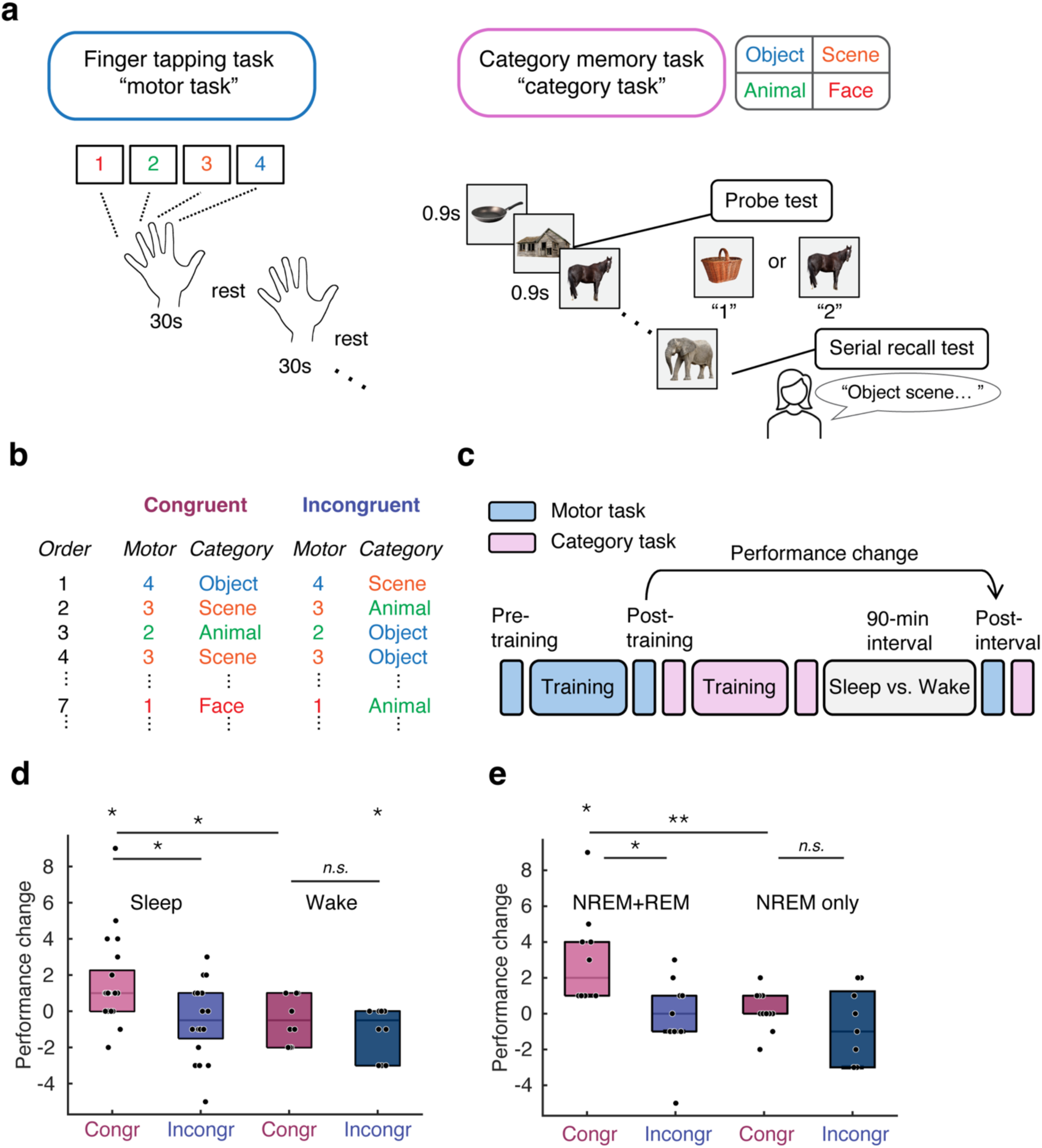
Sleep, but not wakefulness, facilitates declarative-to-procedural memory transfer in the face of a common higher-order rule. **a**, Finger-tapping motor sequence task (motor task, left) and category memory task (category task, right). **b**, Congruent and incongruent conditions for the motor and category tasks. In the congruent condition, the motor and category tasks shared a common higher-order sequence rule, whereas in the incongruent condition, there was no correspondence in the higher-order sequence rule between the two tasks. **c,** Experimental design. After training on the motor and category tasks, participants underwent a 90-min interval with or without sleep, followed by a retest session. Polysomnography was performed during sleep. **d,** Sleep, but not wakefulness, promotes declarative-to-procedural memory transfer. Only participants in the sleep group trained with a shared higher-order sequence rule between the motor and category tasks showed a significant improvement in motor task performance, indicating that declarative-to-procedural memory transfer requires both sleep and a common higher-order rule. Each dot represents the change in the number of correctly typed sequences from the posttraining session to the postinterval test session. The horizontal line inside each box indicates the median, and the lower and upper edges indicate the 25th and 75th percentiles, respectively. Sleep-congruent, *n* = 21; sleep-incongruent, *n* = 20; wake-congruent, *n* = 10; and wake-incongruent, *n* = 10. **e**, Memory transfer manifests after NREM and REM sleep. Only participants in the NREM+REM group trained with a common higher-order sequence rule showed a significant improvement in motor task performance. Box plots indicate the median and quartiles. Each dot represents the change in the number of correctly typed sequences from the posttraining session to the postinterval test session for each participant. NREM+REM-congruent, *n* = 10; NREM+REM-incongruent, *n* = 11; NREM-only-congruent, *n* = 11; NREM-only-incongruent, *n* = 9. \*\**p* < 0.01, \**p* < 0.05, two-tailed Mann‒Whitney U test or Wilcoxon signed-rank test, FDR adjusted. The hand image in Fig. 1a was selected from openclipart.org (https://openclipart.org/detail/147973/hand) and modified. Other images in Fig. 1a were selected from Pixabay (https://pixabay.com/photos/) and modified to resemble the actual task setting.

Participants were randomly assigned to either congruent or incongruent conditions, where a rule was hidden in the congruent condition, inspired by a previous study (*18*). In the congruent condition, the two tasks shared a higher-order common sequence rule, i.e., the sequence in the motor and category tasks matched, whereas the sequences did not match in the incongruent condition (Fig. 1b; see <u>Higher-order sequence</u> in the Materials and Methods section). After participants were trained on the two tasks, a 90-minute interval occurred with (sleep-congruent or sleep-incongruent group) or without sleep (wake-congruent or wake-incongruent group), followed by a retest session (Fig. 1c). Polysomnography was performed during the sleep session (table S1; see PSG measurement in the Materials and Methods section). To examine the behavioral outcomes, the change in the number of correct sequences from posttraining to postinterval in the motor task was measured as the performance change due to the interval (postinterval minus posttraining). Category task performance was measured as the number of correct responses on the probe task and the recall accuracy for the serial recall task. Given sleep’s role in extracting abstract rules (*41*, *42*), we hypothesized that only the sleep-congruent group would show a significant improvement in the motor task.

We indeed found that only the sleep-congruent group showed a significant improvement in the motor task after the interval (Fig. 1d; *n* = 21 participants, one-sample Wilcoxon signed-rank test, two-tailed, *z* = 2.7718, *p* = 0.0224, FDR adjusted). The sleep-incongruent (*n* = 20 participants, one-sample two-tailed Wilcoxon signed-rank test, *z* = −0.6865, *p* = 0.4924, FDR adjusted) and the wake-congruent (*n* = 10 participants, one-sample two-tailed Wilcoxon signed-rank test, *z* = - 1.0433, *p* = 0.3957, FDR adjusted) groups showed no significant difference from their posttraining performance. The wake-incongruent group performed significantly worse after the interval (*n* = 10 participants, one-sample two-tailed Wilcoxon signed-rank test, *z* = −2.0702, *p* = 0.0466, FDR adjusted). The sleep-congruent group performed significantly better than the sleep-incongruent group did (two-tailed Mann‒Whitney U test, *z* = 2.2693, *p* = 0.0363, FDR adjusted), whereas no significant difference in performance was found between the wake-congruent and incongruent groups (two-tailed Mann‒Whitney U test, *z* = 1.0843, *p* = 0.2782, FDR adjusted). The sleep-congruent group performed significantly better than the wake-congruent group did (two-tailed Mann‒Whitney U test, *z* = 2.2540, *p* = 0.0363, FDR adjusted).

The results thus far indicate that participants trained with a common higher-order sequence rule and who had sleep during the interval improved significantly on the motor task after the interval. We next asked whether the category task deteriorated after sleep, given that a previous study showed that memory interaction comes at a cost: while one of the two tasks improves, the other task deteriorates over wakefulness (*18*). However, we found no evidence of memory deterioration for the category task (table S2). No significant difference was found between the groups, and none of the groups showed any change after the interval. These results indicate that sleep, but not wakefulness, promotes declarative to procedural memory transfer without sacrificing declarative memory performance.

Sleepiness (table S3) did not significantly differ between the groups. Therefore, these are unlikely to have influenced the differences in memory transfer between the groups.

### Transfer is masked until REM sleep occurs

Next, to identify the role of REM sleep, participants were classified into two subgroups (see Experimental design in the Materials and Methods section), the NREM+REM and NREM-only groups, for each of the congruent and incongruent conditions. The NREM+REM groups consisted of participants who showed REM after NREM sleep (NREM+REM-congruent, *n* = 10; NREM+REM-incongruent, *n* = 11) during the sleep session, whereas the NREM-only groups consisted of participants who showed only NREM sleep without REM sleep (NREM-only-congruent, *n* = 11; NREM-only-incongruent, *n* = 9).

A significant difference in performance was observed between the NREM+REM-congruent and NREM-only-congruent groups (Fig. 1e; *n* = 21 in total, Mann‒Whitney U test, two-tailed, *z* = 3.0638, *p* = 0.0066, FDR adjusted). The NREM+REM-congruent group performed significantly better than the NREM+REM-incongruent group did (*n* = 21 in total, Mann‒Whitney U test, two-tailed, *z* = 2.6543, *p* = 0.0119, FDR adjusted). No significant difference was observed between the NREM-only-congruent and NREM-only-incongruent groups (*n* = 20 in total, Mann‒Whitney U test, two-tailed, *z* = 1.0087, *p* = 0.3131, FDR adjusted). Only the NREM+REM-congruent group showed a significant improvement after sleep (n = 10, one-sample Wilcoxon signed-rank test, two-tailed, *z* = 2.8421, *p* = 0.0180, FDR adjusted). None of the NREM+REM-incongruent (*n* = 11, one-sample Wilcoxon signed-rank test, two-tailed, *z* = 0.5407, *p* = 0.7849, FDR adjusted), NREM-only-congruent (*n* = 11, one-sample Wilcoxon signed-rank test, two-tailed, *z* = 0.1588, *p* = 0.8738, FDR adjusted), or NREM-only-incongruent groups (*n* = 9, one-sample Wilcoxon signed-rank test, two-tailed, *z* = −1.2036, *p* = 0.4574, FDR adjusted) showed a significant difference from 0. Thus, the motor task improved significantly only in the NREM+REM group in the congruent condition after sleep. These results demonstrate that memory transfer manifests after NREM and REM sleep, leveraging common higher-order information between the declarative and procedural tasks.

### NREM sleep supports processing in local motor circuits for future transfer

We next investigated what neural underpinnings during sleep are involved in memory transfer. With respect to NREM sleep, slow wave–spindle coupling has been proposed as a key driver of memory processing (*16*, *43*). When sleep spindles are nested in the up state of slow waves, reactivated information is redistributed and stabilized in the neocortex (*44*). The causal effects of slow wave–spindle coupling on memory formation have been reported in multiple studies (*43*, *45*). If slow wave–spindle coupling drives multiregional communication across regions (*46*, *47*) for memory transfer, we should observe a significant difference in coherence between the conditions. To this end, we performed EEG source reconstruction (see EEG source reconstruction in the Materials and Methods section) and selected four regions of interest (ROIs) as potential waypoints for memory transfer, namely, the SMA, superior parietal cortex, hippocampus, and mPFC (fig. S1; Regions of interest (ROIs) in the Materials and Methods section) on the basis of earlier studies (*13*, *48–55*). Among the possible pairs, five pairs (white arrows), i.e. mPFC–hippocampus (left and right) (*54–56*), right SMA–parietal (*53*), right SMA–hippocampus (left and right) (*48–50*) were included for coherence measurement based on prior studies (see Regions of Interests and Coherence analysis in the Materials and Methods section). To test whether multiregional communication occurs in association with slow wave–spindle coupling, slow-wave events were classified into ‘coupled’ and ‘uncoupled’ events (fig. S2, Sleep event detection in the Materials and Methods section), and coherence values for 0.5–2 Hz corresponding to the slow-wave band were measured for each of the events, averaged for each participant, and compared across the conditions.

Compared with the incongruent condition, the congruent condition was associated with significantly *decreased* connectivity between the mPFC and the left (Fig. 2a; *n* = 23 in total, Mann‒Whitney U test, two-tailed, *z* = −2.3695, *p* = 0.0445, FDR adjusted) and right hippocampus (*n* = 23, Mann‒Whitney U test, two-tailed, *z* = −2.6772, *p* = 0.037, FDR adjusted) (Fig. 2a, c, d). Compared with the incongruent condition, the right SMA showed moderately *increased* connectivity between the parietal cortex for the congruent condition during slow-wave sleep (Fig. 2e, *n* = 23, Mann‒Whitney U test, two-tailed, *z* = 2.0618, *p* = 0.0653, FDR adjusted). No significant difference was found in the other two ROI pairs (Fig. 2f, g; right SMA–left hippocampus, *n* = 23, Mann‒Whitney U test, two-tailed, *z* = −1.2617, *p* = 0.2071, FDR adjusted; right SMA–right hippocampus, *n* = 23, Mann‒Whitney U test, two-tailed, *z* = 2.0618, *p* = 0.2071, FDR adjusted).

**Fig. 2:**
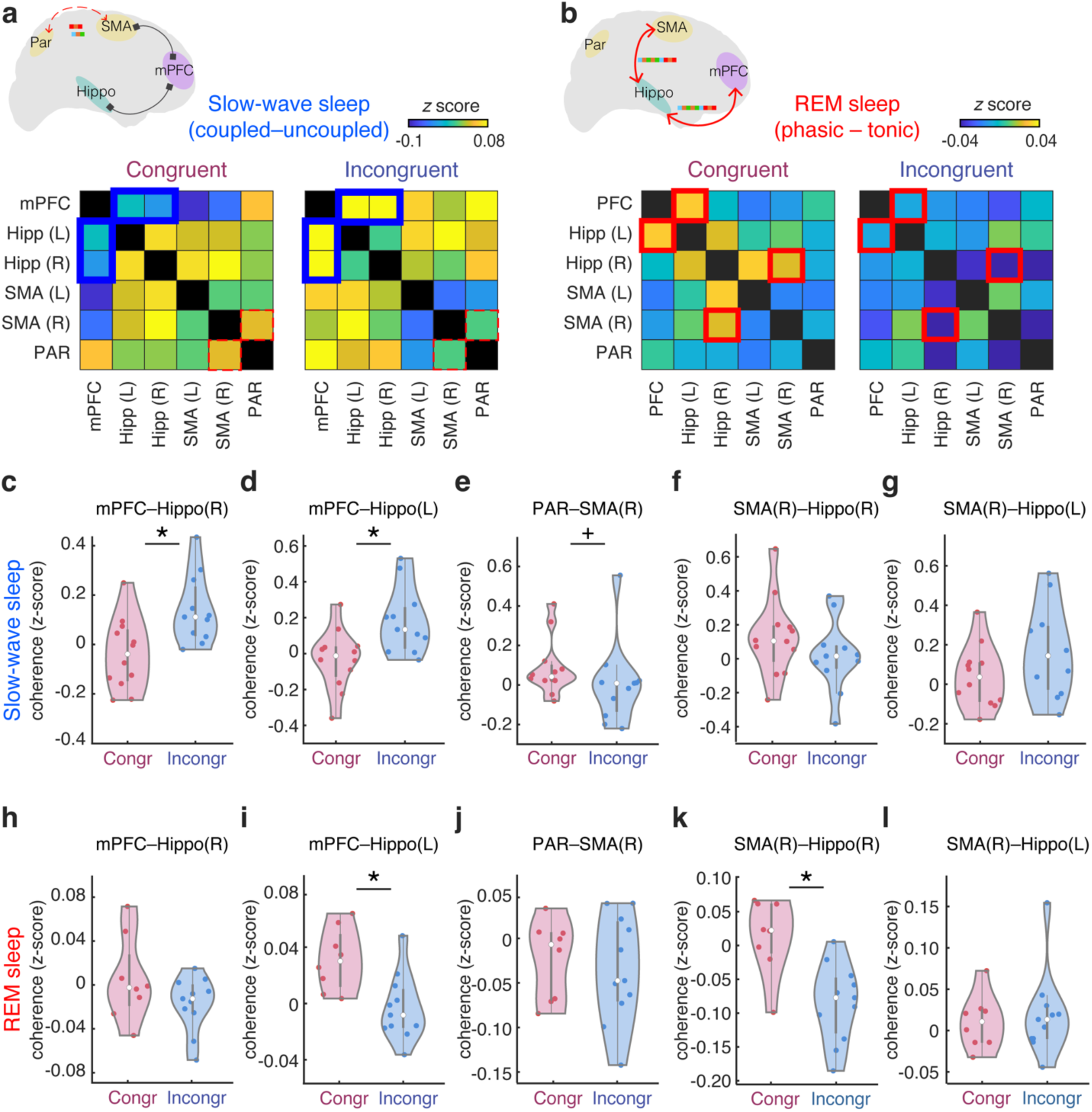
Disconnected and interconnected network during NREM and REM sleep. **a,** Coherence matrix of the slow wave band (0.5–2 Hz) during NREM (slow-wave) sleep in the congruent (left) and incongruent (right) conditions. The color scales indicate the range of z-transformed coherence values. The cells with the blue line indicate a significant difference between the congruent and incongruent conditions, corresponding to **c** and **d** below. The brain image shows where a significant or moderate difference was found in the coherence values between the conditions (red, stronger in the congruent; gray, weaker in the congruent). **b,** Coherence matrix of the theta band (5–9 Hz) during REM phasic sleep in the congruent (left) and incongruent (right) conditions. The color scales indicate the range of z-transformed coherence values. The cells with the red line indicate a significant difference between the congruent and incongruent conditions, corresponding to **i** and **k** below. The brain image shows where a significant difference was found in the coherence values between the conditions (red, stronger in the congruent). **c-g**, Violin plots showing coherence values during slow-wave sleep for the pairs of interest. Coherence values between the mPFC and the right (**c**) and left (**d**) hippocampus significantly differed between the congruent and incongruent conditions (*p* < 0.05). There was a trend in the difference in the coherence values between the parietal region and the SMA (**e**). No significant difference was found between the conditions for the right SMA and the hippocampus (**f** and **g**). Congruent condition, *n* = 12. Incongruent condition, *n* = 11. **h-l**, Connectivity during REM sleep for the pairs of interests. Coherence values between the mPFC and the left hippocampus (**i**) and coherence values between the SMA and the right hippocampus (**k**) were significantly different between the congruent and incongruent conditions (*p* < 0.05). No significant difference was found between the conditions for the other pairs (**h**, **j**, and **l**). Violin plots show medians and quartiles. Congruent condition, *n* = 12. Incongruent condition, *n* = 11. \**p* < 0.05, ^+^*p* < 0.1, two-tailed Mann‒Whitney U test, FDR adjusted.

We found that these differences in connectivity were specific to slow-wave sleep. No significant differences between the conditions were found in stage N2 connectivity (*n* = 25 in total, right SMA–parietal: Mann‒Whitney U test, *z* = 0.3011, *p* = 0.9346, FDR adjusted; right SMA–left hippocampus: Mann‒Whitney U test, two-tailed, *z* = 0.0821, *p* = 0.9346, FDR adjusted; right SMA–right hippocampus: Mann‒Whitney U test, two-tailed, *z* = 0.3011, *p* = 0.9346, FDR adjusted; mPFC–left hippocampus: Mann‒Whitney U test, two-tailed, *z* = 0.1916, *p* = 0.9346, FDR adjusted; mPFC–right hippocampus: Mann‒Whitney U test, two-tailed, *z* = −0.4653, *p* = 0.9346, FDR adjusted). These results suggest that during deep NREM sleep, when slow waves and sleep spindles are abundant in synchrony, motor memory is processed in a local motor circuit, possibly to strengthen motor memory, while keeping other regions disconnected, utilizing higher-order information.

To investigate whether the differences in functional connectivity are due to local increases in these activities, we measured the density of slow waves, sleep spindles, and coupling events for each ROI (Density measurement in the Materials and Methods section). We found no significant differences in any of the ROIs for slow wave density (fig. S3; two-tailed independent-samples *t* test, Congruent *n* = 12, Incongruent *n* = 11, *t*_21_= −1.7432, *p* = 0.0959), sleep spindle density (fig. S4; two-tailed independent-samples *t* test, Congruent *n* = 12, Incongruent *n* = 11, *t_21_*= 0.0127, *p* = 0.9900), or coupling density (fig. S5; two-tailed independent-samples *t* test, Congruent *n* = 12, Incongruent *n* = 11, *t_21_*= −1.4684, *p* = 0.1568). Thus, connectivity between regions, rather than local activity strength, is likely involved in memory transfer.

### REM sleep breaks the procedural–declarative system boundaries, allowing transfer

We next asked whether connectivity between the brain regions was significantly different between the congruent and incongruent conditions during REM sleep. REM sleep has been classified into phasic and tonic periods (see REM phasic and tonic periods in the Materials and Methods section) (*17*, *57–59*). The phasic period is marked by conspicuous physiological activities, including rapid eye movements. The tonic period is defined as the suppression of these events. While the phasic period is involved in memory processing, external stimuli are processed in a wake-like manner during the tonic period (*57–59*). Thus, to study the role of REM sleep in memory transfer, we investigated how the brain is modulated during the phasic period compared with the tonic period. After rapid eye movements were detected semiautomatically, PSG data corresponding to REM sleep were classified into phasic or tonic periods every 3 s. REM sleep epochs containing at least two rapid eye movements were classified as phasic periods, and those having fewer than two rapid eye movements (one or none) were classified as tonic periods (fig. S6; see Sleep event detection in the Materials and Methods section). Then, coherence values for 5–9 Hz, corresponding to the theta band, causally implicated in memory processing (*27*), were measured for each 3-s epoch. The same five ROI pairs as in NREM sleep were selected for the measurement of coherence.

In contrast to during NREM sleep, the congruent condition was associated with an overall increase in across-area connectivity during REM sleep (Fig. 2b). Compared with the incongruent condition, the congruent condition significantly *increased* functional connectivity between the right SMA and the right hippocampus (Fig. 2k; *n* = 19 in total, Mann‒Whitney U test, two-tailed, *z* = 2.8487, *p* = 0.01825, FDR adjusted) and between the mPFC and the left hippocampus (Fig. 2i; Mann‒Whitney U test, two-tailed, *z* = 2.6836, *p* = 0.01825, FDR adjusted). No other ROI pairs showed a significant difference between the conditions (Fig. 2h, j, l; right SMA–parietal: Mann‒Whitney U test, two-tailed, *z* = 0, *p* = 1.00; right SMA–left hippocampus: Mann‒Whitney U test, two-tailed, *z* = −0.2064, *p* = 1.00; mPFC–right hippocampus: Mann‒Whitney U test, two-tailed, *z* = 1.1147, *p* = 0.4417; all FDR adjusted).

These results suggest that during REM sleep, the extraction of regularities, abstraction, and knowledge generation occur through changes in interactions among the SMA, the hippocampus, and the mPFC.

To test the possibility that the differences in coherence between the conditions were caused simply by differences in the number of phasic periods, we measured REM phasic density (the number of phasic periods/all REM epochs x 100). There was no significant difference in REM phasic density between the conditions (two-tailed independent-samples t test, Congruent *n* = 8, Incongruent *n* = 11, *t*17= −0.7560, *p* = 0.4600). Thus, the differences in coherence values cannot be attributed to changes in the number of phasic events.

### Memory state becomes instantaneously unstable during NREM sleep, as revealed by the time-resolved MRS-based E/I ratio

While it has been assumed that unstable memory facilitates memory transfer during wakefulness (*18*), we instead hypothesized that memory becomes hyperstable during REM sleep (*23*) to tolerate across-system transfer once it has been destabilized for modifications during NREM sleep. If so, shifts in the memory state should occur in a close relationship with sleep microstructures—slow wave–spindle coupling and REM phasic periods—when changes in functional connectivity take place. To address this question, we measured MRS-based excitation-to-inhibition ratios (E/I ratios) at the system level E/I during sleep as a readout for memory stability (*19*, *23*, *34*).

A challenge in MRS is its poor temporal resolution (*35*). The temporal resolution has been limited to 2–10 min on 3-T MRI. However, sleep microstructures can transiently emerge and then subside within a few seconds or less (*37*, *38*). If these sleep microstructures actively drive memory transfer, averaging the signals could falsely obscure any effects that are specific to these states. Thus, E/I ratios should be measured at much higher temporal resolution to investigate changes in memory stability in association with microstructures. Thus, in experiment 2, we developed a new simultaneous MRS and PSG method, leveraging ultrahigh-field 7-Tesla MRI to increase temporal resolution (Fig. 3a; see Experimental design and MRI acquisition in the Materials and Methods section). We succeeded for the first time in estimating time-varying E/I ratio changes during NREM and REM sleep (Fig. 3b & c).

**Fig. 3:**
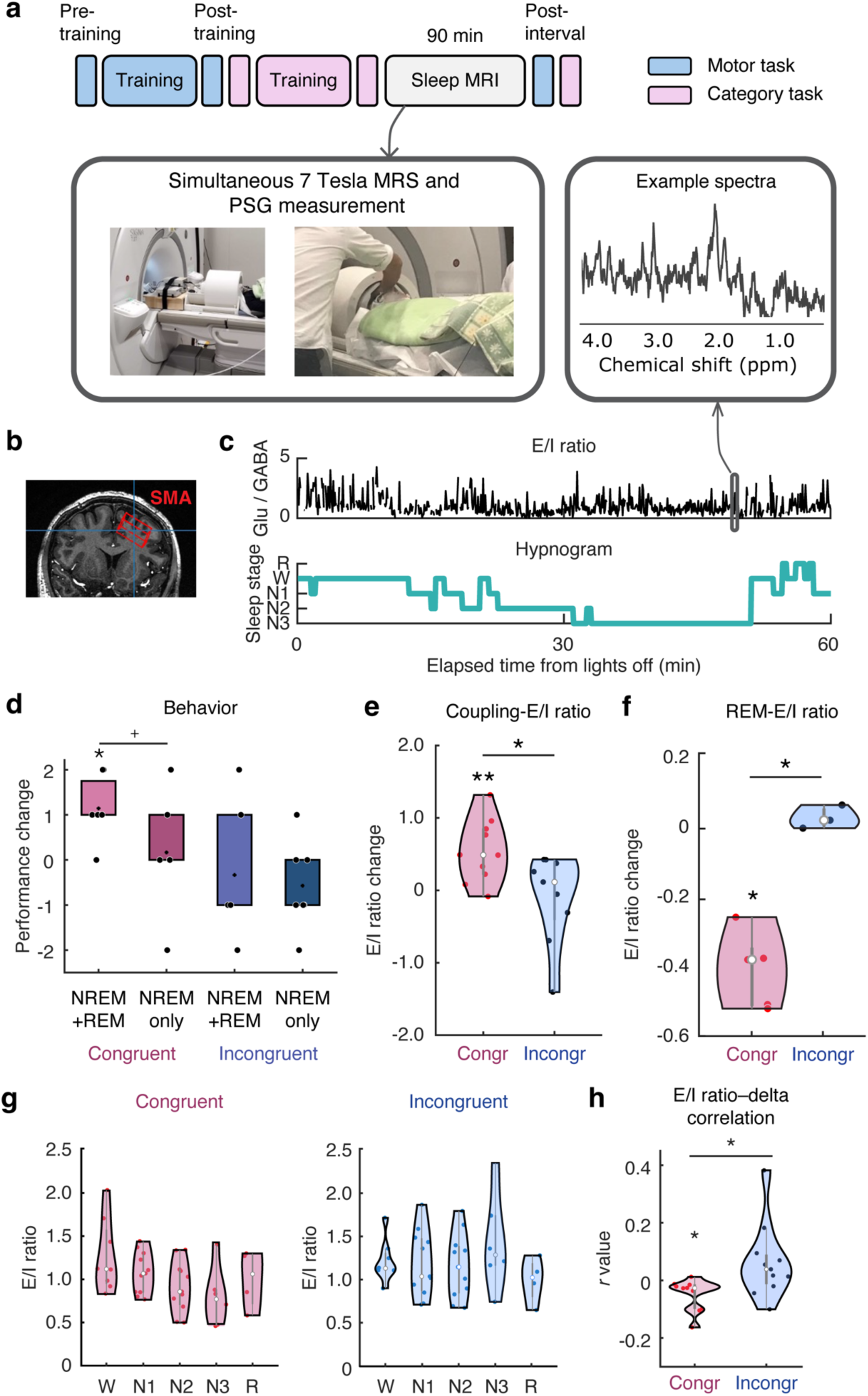
Time-resolved E/I ratio changes during non-REM and REM sleep. **a,** Experimental design. As in experiment 1, after training on the motor and category tasks, a 90-min sleep session occurred, followed by retesting. MRS was measured simultaneously with PSG during sleep (inset). **b**, Volume placement in the SMA for MRS measurement. **c**, Example 5-sec resolution E/I ratio changes (top) and the corresponding hypnogram (bottom) of a representative dataset. An inset above shows example spectra measured from a single segment. Some parts could not be fitted because of the low signal-to-noise ratio. The E/I ratios were measured according to the glutamate and GABA concentrations. **d**, Behavioral results of experiment 2, replicating the results of experiment 1. Only the NREM+REM group in the congruent condition showed a significant improvement in the motor task (Wilcoxon signed-rank test, *n* = 7, two-tailed, *z* = 2.2711; *p* = 0.0231). The NREM-only group in the congruent condition (Wilcoxon signed-rank test, two-tailed, *n* = 6, *z* = 0.2722, *p* = 0.7855) and the incongruent condition (NREM+REM, *n* = 6, *z* = −0.5407, *p* = 0.5887; NREM-only, *n* = 7, *z* = −1.4142, *p* = 0.1573) showed no improvement after sleep. **e**, The E/I ratio increases in association with slow wave–spindle coupling only in the congruent condition. Congruent, *n* = 10; incongruent, *n* = 10. **f**, The E/I ratio decreases in association with the REM phasic period only in the congruent condition. **g**, Sleep stage–dependent E/I ratio for the congruent (left) and incongruent (right) conditions. **h**, Correlations between the E/I ratio and delta power (0.5–4 Hz) during NREM sleep. \*\**p* < 0.01, \**p* < 0.05, ^+^*p* < 0.1, Mann‒Whitney U test or Wilcoxon signed-rank test (two-tailed, FDR corrected).

We trained participants on the motor and category tasks before sleep and retested them on the same tasks after sleep as in experiment 1 (Fig. 3a). After the participants were trained on the two tasks, they had a 90-minute nap inside an MRI scanner (table S4), followed by a retest session. Inside the MRI scanner, simultaneous MRS and PSG measurements were performed to measure the MRS-based E/I ratio changes from the right SMA (Fig. 3b & c; MRI acquisition in the Materials and Methods section). The change in the number of correct sequences from posttraining to postinterval in the motor task was measured as the performance change due to the interval (postinterval minus posttraining), as in experiment 1.

The performance changes replicated the results of experiment 1 (Fig. 3d): Only the participants who had REM after NREM sleep and who were trained with a common higher-order sequence rule (NREM+REM-congruent) showed a significant improvement in the motor task (*n* = 7, two-tailed one-sample Wilcoxon signed-rank test against 0, *z* = 2.2711, *p* = 0.0462), while the incongruent condition showed no significant difference (n = 6, two-tailed one-sample Wilcoxon signed-rank test against 0, *z* = −0.5407, *p* = 0.5887). The NREM+REM-congruent group performed better than the incongruent group did, although the difference was only marginally significant (*n* = 13 in total, two-tailed Mann‒Whitney U test, *z* = 1.7099, *p* = 0.0873). No significant difference was observed between the congruent and incongruent conditions in the NREM-only group (*n* = 13 in total, two-tailed Mann‒Whitney U test, *z* = 1.1119, *p* = 0.2662). None of the groups showed any change after the interval for the category task (table S5). Thus, the transfer of a higher-order representation was effective only after NREM and REM sleep, replicating the results of experiment 1.

We next tested whether the memory state becomes unstable for transfer during NREM sleep. We classified NREM sleep into coupled and uncoupled slow-wave epochs on the basis of NREM sleep events (Sleep event detection in the Materials and Methods section) every 5 s and coregistered them with MRS data that were measured every 5 s. E/I ratios were calculated by taking the glutamate to GABA concentration ratios for each 5-second epoch (fMRS analysis for sleep microstructures in the Materials and Methods section). Afterward, the E/I ratios during NREM sleep were averaged for each of the coupled and uncoupled epochs, and the uncoupled E/I ratios were subtracted from the coupled E/I ratios to obtain the coupling-specific E/I ratio change (see Inclusion and exclusion of data in the Materials and Methods section).

Does the memory state become temporarily unstable when it goes through processing in a local circuit, as has been previously reported in other types of learning (*19*, *32*)? If this is the case, the E/I ratio should become more excitatory in association with slow wave–spindle coupling when local motor memory processing takes place. The coupling-specific E/I ratio in the congruent condition was indeed significantly *greater* than that in the incongruent condition (Fig. 3e; two-tailed independent-samples *t* test between conditions, *n* = 19, *t*_19_ = 2.6335, *p* = 0.0174, 95% *CI* = [0.1269, 1.1492]). Only the coupling-specific E/I ratio in the congruent condition showed a significant difference from 0 (two-tailed one-sample *t* test against 0, *n* = 10, *t*_9_ = 3.9679, *p* = 0.0033, 95% *CI* = [0.2329, 0.8507]), whereas the E/I ratio in the incongruent condition showed no significant difference from 0 (two-tailed one-sample *t* test against 0, *n* = 9, *t*_8_ = −0.4677, *p* = 0.6525). Thus, the memory state becomes instantaneously unstable in association with slow wave–sleep spindle coupling during NREM sleep only when participants are trained on two tasks that share common higher-order information.

### Memory state becomes hyperstable for transfer during phasic REM sleep

We next investigated whether the memory state becomes highly stable when it is transferred beyond a single memory system. If this is the case, the E/I ratio should become more inhibitory in association with the REM phasic period when communication among brain regions occurs.

REM sleep was classified into phasic and tonic periods in accordance with rapid eye movements (fMRS analysis for sleep microstructures in the Materials and Methods section) every 5-s epoch. The E/I ratios during REM sleep were averaged for each of the phasic and tonic periods, after which the tonic E/I ratios were subtracted from the phasic E/I ratios to obtain REM phasic-specific E/I ratio changes. Thus, a higher or lower E/I ratio indicates a more unstable or stable memory state during the phasic period than during the tonic period. Compared with those in the incongruent condition, the REM phasic-specific E/I ratios in the congruent condition were significantly *lower* (Fig. 3f; *n* = 8 in total, two-tailed Mann‒Whitney U test, *z* = −2.0870, *p* = 0.0369). Only the congruent condition showed a significant difference from 0 (*n* = 5, two-tailed one-sampled Wilcoxon signed-rank test against 0, *z* = −2.0226, *p* = 0.0431), while no significant difference was found in the incongruent condition (*n* = 3, two-tailed one-sampled Wilcoxon signed-rank test against 0, *z* = 0.5345, *p* = 0.5930). Thus, the memory states become rapidly hyperstable in association with REM phasic periods when memory transfer occurs.

One may wonder whether the differences in E/I ratios between the congruent and incongruent conditions were due to differences in MRS data quality. We confirmed that there was no significant difference in MRS data quality measures between the congruent and incongruent conditions (table S6; see MRS data quality in the Materials and Methods section). We also confirmed that sleepiness (table S7) did not significantly differ between the groups. Therefore, these cannot be attributed to the differences in E/I ratios between the groups.

### Memory state gradually stabilizes as sleep deepens

The results thus far demonstrate that memory state shifts are strongly linked to sleep microstructures during NREM and REM sleep. We further asked whether memory stability changes with sleep depth, given previous findings reporting sleep stage–based E/I ratio changes in visual perceptual learning (*23*). To address this, we measured the E/I ratios every 30 s, averaged them for each sleep stage and tested whether they were significantly different from wakefulness (fMRS analysis for sleep stages in the Materials and Methods section).

E/I ratios were significantly different across sleep stages for the congruent condition (Kruskal‒Wallis test, *H*_4_ = 11.36, *p* = 0.0228), whereas no significant difference was found for the incongruent condition (Kruskal‒Wallis test, *H*_4_ = 3.69, *p* = 0.4492). We next investigated whether the differences in the E/I ratios among sleep stages in the congruent condition were associated with sleep depth. Sleep depth is correlated with delta-band power (*60*, *61*). Thus, if the sleep-stage-dependent E/I ratio decrease is indeed associated with sleep depth, E/I ratio changes should correlate with delta power, and the correlation should be stronger in the congruent condition than in the incongruent condition. Multitaper analysis (*62*, *63*) was performed on EEG data from the right SMA, where the E/I ratio was measured every 30 s. Then, correlation coefficients between delta power during NREM sleep and E/I ratios were measured (Correlation analysis between the E/I ratio and delta power in the Materials and Methods section). There was a significant difference in the correlation coefficient between the congruent and incongruent conditions (Fig. 3h; two-tailed independent samples *t* test, *n* = 21, *t*_19_ = −2.1438, *p* = 0.04524, 95% *CI* = [-0.1458, −0.0017]). Only the congruent condition showed a significant negative correlation between delta power and the E/I ratio (two-tailed one sample *t* test against 0; congruent: *n* = 10, *t*_9_ = −2.8643, *p* = 0.0186, 95% *CI* = [-0.0882, −0.0104]; incongruent: *n* = 11, *t*_10_ = 1.5999, *p* = 0.1407, 95% *CI* = [-0.0245, 0.1494]). The inclusion of all the time course data, which was not limited to NREM sleep, yielded similar results (fig. S7). These findings indicate that the memory state gradually becomes stable as sleep deepens.

### Different neurochemical states subserve memory state shifts during NREM and REM sleep

The neurochemical basis of memory stability could differ depending on sleep stage and microstructure, involving excitatory and/or inhibitory mechanisms (*64*, *65*). Thus, we next investigated whether the glutamate or GABA concentration could account for the changes in the E/I ratio during sleep.

First, we found that the change in the E/I ratio for slow wave–spindle coupling was driven by the change in glutamate concentration (Fig. 4a). The glutamate concentration was significantly greater in the congruent condition than in the incongruent condition (two-tailed independent samples *t* test, *n* = 19 in total, *t*_17_ = 3.1141, *p* = 0.0063, 95% *CI* = [1.3449, 6.9953]). The glutamate concentration in the congruent condition was significantly different from 0 (two-tailed one-sample t test against 0, *n* = 10, *t*_9_ = 2.7918, *p* = 0.0210), whereas no significant difference was found for the incongruent condition (two-tailed one-sample *t* test against 0, *n* = 9, *t*_8_ = −1.5648, *p* = 0.1563). No significant difference in the GABA concentration was detected between the conditions (two-tailed independent-samples *t* test, *n* = 19 in total, *t*_17_ = −1.0814, *p* = 0.2946, 95% *CI* = [-8.0027 2.5791]). The GABA concentration did not significantly differ from 0 for either the congruent (two-tailed one-sample *t* test against 0, *n* = 10, *t*_9_ = −0.8680, *p* = 0.4080) or incongruent condition (two-tailed one-sample *t* test against 0, *n* = 9, *t*_8_ = 0.7354, *p* = 0.4831).

**Fig. 4:**
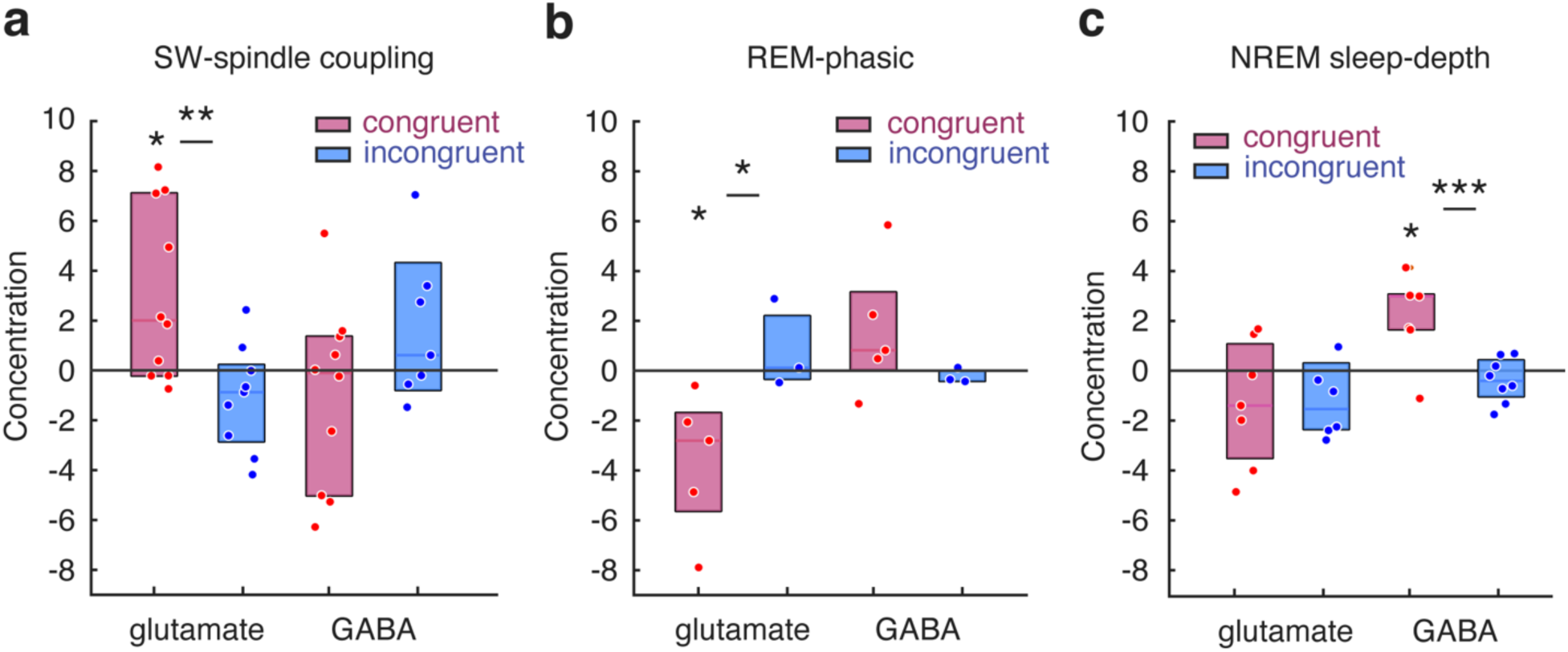
Sleep microstructures and depth-related neurochemical changes. **a,** Slow wave–spindle coupling-specific glutamate and GABA concentrations. The glutamate concentration associated with slow wave–spindle coupling was significantly greater in the congruent condition than in the incongruent condition. The glutamate concentration in the congruent condition was significantly different from 0, whereas no significant difference was detected in the incongruent condition. No significant difference in the GABA concentration was detected between the conditions. The GABA concentration did not significantly differ from 0 for either the congruent or incongruent condition. SW, slow wave. **b,** REM phasic-associated glutamate and GABA concentrations. The REM phasic glutamate concentration was significantly lower in the congruent condition than in the incongruent condition. The REM phasic-specific concentration level in the congruent condition was significantly different from 0, whereas no significant difference was found in the incongruent condition. No significant difference in the GABA concentration was detected between the conditions or baseline. **c,** NREM sleep-depth-dependent changes in GABA and glutamate concentrations. The concentration of GABA during NREM sleep was significantly greater than that during wakefulness in the congruent condition and significantly greater than that during NREM sleep in the incongruent condition. No significant difference was found for the incongruent condition. The glutamate concentration did not significantly differ during NREM sleep between the congruent and incongruent conditions, and there was no significant difference between the conditions. * *p*< 0.05, ^+^*p* < 0.1, two-tailed Mann‒Whitney U test, Wilcoxon signed-rank test, independent-samples or one-sample *t* tests.

Next, we found that the REM-phasic glutamate concentration was significantly lower in the congruent condition than in the incongruent condition (Fig. 4b; *n* = 8 in total, two-tailed Mann‒Whitney U test, *z* = −2.0870, *p* = 0.0369). The REM phasic-specific concentration level in the congruent condition was significantly different from 0 (*n* = 5, two-tailed one-sample Wilcoxon signed-rank test against 0, *z* = −2.0226, *p* = 0.0431), whereas no significant difference was found for the incongruent condition (*n* = 3, two-tailed one-sample Wilcoxon signed-rank test against 0, *z* = −1.0690, *p* = 0.2850). No significant difference in the GABA concentration between the conditions or baseline was detected (congruent vs. incongruent: *n* = 8 in total, two-tailed Mann‒Whitney U test, *z* = 0.2981, *p* = 0.7656; congruent against 0: *n* = 5, two-tailed one-sampled Wilcoxon signed-rank test, *z* = −0.4045, *p* = 0.6858; incongruent against 0: *n* = 3, two-tailed one-sampled Wilcoxon signed-rank test, *z* = −1.0690, *p* = 0.2850).

Finally, we tested whether sleep stage–dependent changes in the E/I ratio were driven by changes in the concentration of glutamate or GABA. To this end, we measured changes in glutamate and GABA concentrations during NREM sleep normalized by wakefulness (calculated as the mean of the N2 and N3 sleep stages minus the mean during wakefulness; Fig. 4c). We averaged the N2 and N3 stages because sleep stage–dependent changes in the E/I ratio occurred mainly during stable NREM sleep (Fig. 3g). The GABA concentration during NREM sleep was significantly greater than that during wakefulness in the congruent condition (two-tailed one-sample *t* test against 0, *n* = 7, *t*_6_ = 3.3868, *p* = 0.0147, 95% *CI* = [0.6035, 3.7457]) and significantly greater than that in the incongruent condition (two-tailed independent-samples t test, *n* = 15, *t*_13_ = 3.7865, *p* = 0.0023, 95% *CI* = [1.1148 4.0768]). No significant difference from baseline was found for the incongruent condition (two-tailed one-sample *t* test against 0, *n* = 8, *t*_7_ = −1.3447, *p* = 0.2206). The glutamate concentration did not significantly differ during NREM sleep in the congruent (two-tailed one-sample t test against 0, *n* = 7, *t*_6_ = −1.3872, *p* = 0.2147) or incongruent condition (two-tailed one-sample t test against 0, *n* = 8, *t*_7_ = 0.0539, *p* = 0.9585), and there was no significant difference between the conditions (two-tailed independent-samples *t* test, *n* = 15, *t*_13_ = −0.7379, *p* = 0.4737).

Thus, instantaneous shifts in the E/I ratio during sleep, whether associated with slow wave–spindle coupling or the REM phasic period, are driven mainly by changes in the concentration of glutamate, whereas gradual sleep stage–dependent changes are driven by changes in the concentration of GABA. These results suggest that distinct neural underpinnings are involved in changes in memory stability during sleep.

## Conclusions

The results of this study demonstrate that sleep supports memory transfer using higher-order sequence information. During NREM sleep, the memory state becomes temporarily unstable in the SMA while functional connectivity between the hippocampus and mPFC decreases in association with slow wave–spindle coupling. Conversely, during REM sleep, the memory state is hyperstabilized, while the SMA, the hippocampus, and the mPFC are interconnected. In addition to these abrupt memory state shifts, sleep depth–dependent gradual stabilization occurs. On the basis of these findings, we propose that memory transfer occurs in multiple phases throughout sleep (Fig. 5). The first stage occurs during NREM sleep, in which unstable memory traces undergo further modification within local circuits, driven by the coupling of brain oscillatory activities. The subsequent stage that follows local processing manifests specifically during REM sleep, during which multiple brain networks are interconnected while memory is hyperstabilized in association with abstraction, resulting in transfer.

**Fig. 5:**
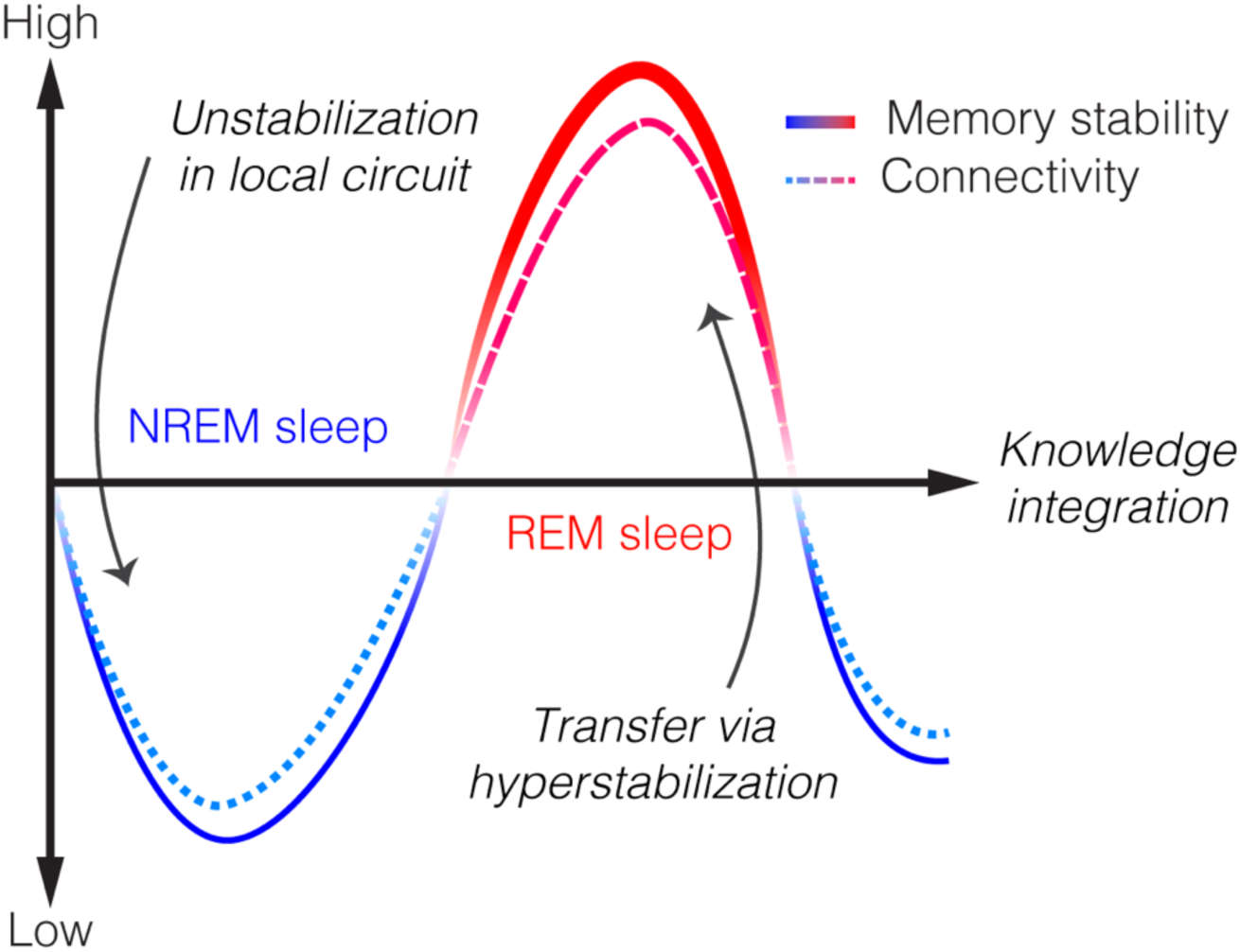
Hypothetical process for memory transfer and knowledge integration. Memory transfer occurs in at least two stages: local and interconnected processing through shifts in memory stability. During the first stage (blue line), memory is so unstable that it is further modified in a local circuit during NREM sleep in association with slow wave–spindle coupling. The next stage manifests during REM sleep (red line) when brain regions are interconnected. This connection allows memory to be transferred beyond a single memory system for the extraction of rules and abstraction to create new knowledge. The memory state is highly stable at this stage. These dynamic, recurrent network modulations and shifts in memory stability during sleep support adaptive and innovative behaviors in humans. While this hypothesis is based on a prior model of visual learning (*23*), unlike the previous model, the current hypothesis is not limited to a single task or learning.

We further demonstrate that rapid and slower shifts in memory state are modulated by different neurochemical conditions. Rapid memory state shifts are driven by increases (NREM sleep) or decreases (REM sleep) in glutamate, while GABA modulates the slower sleep-stage associated E/I ratio decrease in correlation with delta activity. The rapid modulation linked to oscillation nesting and an REM-substate (phasic periods) may reflect cellular excitability modulations with cortical up and down states that could be directly associated with memory transformation. It is reported that Ca^2+^ activity in pyramidal neurons is augmented when spindles are nested in slow waves (*16*, *66*), which are significantly reduced during REM sleep (*67*). These biphasic changes in neuronal activities may support flexible memory reformation and strengthening while the brain is largely offline. The sleep-depth-dependent slower stabilization may represent synaptic renormalization, down-regulating the net synaptic potentiation that arises during training (*68*, *69*). The slower sleep-depth dependent E/I ratio decrease and GABA-delta correlation during NREM sleep align well with the previous findings regarding down-regulation during slow-wave sleep (*21*, *70*). These results directly show evidence that memory states can go through both unstable and stable states with the latter associated with sleep depths which resolves long-standing controversies regarding learning and memory.

This is not the first study to propose sequential roles played by NREM and REM sleep. The sequential hypothesis (*71*, *72*) posits that slow-wave sleep distinguishes memories from irrelevant or competing traces that undergo downgrading or elimination. The processed memories are then integrated with existing memories during REM sleep (*71*, *73*). The complementary processing during NREM and REM sleep has also been reported in visual learning, with the former associated with increased E/I ratios and the latter associated with decreased E/I ratios (*23*). The present findings strengthen the view that NREM and REM sleep sequentially process memory by clarifying how memories are modified and transferred during each of these stages. During NREM sleep, it is possible that while chunked information is sent from the parietal region to SMA, other potentially interfering information is suppressed, thereby strengthening motor memory. During REM sleep, the memory state was hyperstabilized when the SMA–hippocampus and hippocampus–mPFC were connected, enabling memory transfer. Thus, the role of REM sleep may be to transform memory into a robust entity, or schematizing, while making brain regions more communicable, resulting in the integration of memories. Overall, these processes could be the underlying mechanism of knowledge integration that occurs specifically during sleep.

The present findings do not contradict the consolidation models, including the active systems consolidation model, which elegantly elucidates the role of slow-wave sleep in memory (*22*, *74*). We speculate that local motor processing during slow-wave sleep (Fig. 2a) is closely related to the consolidation process reported in previous studies (*22*). Our results revealed that connectivity between the SMA and the parietal region moderately increased, whereas the E/I ratio increased in the SMA, suggesting motor network-level reactivation during slow wave–sleep spindle coupling (*75*). Moreover, we found that the hippocampus–mPFC connectivity decreased concurrently. These findings may resolve contradictions regarding cross-area communication during slow-wave sleep being regional when the hypothesized mechanisms of systems consolidation are global. That is, the brain network could be indeed globally regulated (*22*) but works in a way that each circuit works independently to successfully modulate local memory entities during slow-wave sleep (*49*).

The behavioral consequences of memory interactions could differ considerably between wakefulness and sleep. A previous study elegantly revealed that memory instability is one of the requisites for motor and word-list memory interactions during wakefulness (*18*). Sequential training on motor and word-list tasks was followed by their interactions when common elements existed between the tasks, as in our study. However, the interaction of the two memories resulted in the maintenance of only one of the two tasks during wakefulness. While a previous study suggested that memory instability is key for such memory interference, our results show that memory is highly stable during REM sleep, resulting in no decline in either of the tasks. Thus, while memories with the same higher-order regularities are prioritized, whether the interactions result in facilitation of memories depends on whether sleep (REM sleep) occurred after training because of differences in neuromodulatory tones (*76*) and plasticity mechanisms associated with brain oscillations (*13*, *16*, *66*).

Despite the inherent limitations of MRS as a noninvasive technique, the present study has for the first time quantified memory modulations driven by sleep microstructures in humans, by developing a simultaneous MRS and PSG setup in a 7-Tesla MRI environment. This method will open up new opportunities for investigating neurochemical changes in association with brain oscillatory activities, which can be captured only by EEG. The use of this method is not limited to spontaneous activities during sleep. For instance, an examination of how memory stability changes in event-related designs, such as when combined with reactivation methods (targeted memory reactivation) to selectively reactivate and modulate specific memory traces (*28*, *77*, *78*) or investigations in clinical populations (*79*), should be useful.

There are limitations in the present study that should be addressed in future studies. First, we report the MRS-based E/I ratio, which is a system-level E/I ratio, determined from aggregated concentrations of glutamate and GABA measured in a large brain region (7200 mm^3^). While an animal study has shown that MRS-detected levels of glutamate and GABA reflect the overall activity of excitatory and inhibitory neural processes (*80*), it remains unclear which cellular compartment, for example, the extracellular space, synaptic vesicles, the synaptic cleft, or the cytosol, underlies the changes in glutamate and GABA levels estimated here (*81*). Second, MRS-based E/I ratios are not the same as EEG-based E/I ratios (*82*, *83*), although these ratios are correlated (*84*). Although the comparison between MRS-based and EEG-based E/I ratios is not within the scope of the current study, it would be an interesting topic to pursue in the future.

Finally, our data were obtained from afternoon naps, and the sleep sessions were restricted to 90 minutes. Thus, how processing during NREM and REM sleep changes during nightly sleep, where sleep cycles repeat, is unclear. Despite these limitations, the results of the present study suggest the following important possibility: a well-consolidated NREM–REM sleep cycle may provide lingering processes for knowledge generation, supporting adaptive and innovative behaviors in humans.

## Materials and Methods

### Participants

To estimate the number of participants required to reliably indicate the effect of a daytime nap on skill learning, we applied the G*Power program (*85*) with the power set at 0.8 and the required significance α-level at 0.05, two-tailed, to previously published behavioral data (*86*). While no studies have tested the effects of sleep on memory transfer using finger tapping motor skill learning before the current study, motor and perceptual skills have common characteristics (*24*). Thus, we have adopted the same criterion for the present study. Improvements in performance after sleep were used as the dependent variable, and a *t* test was used as the test family for the G*Power program. The results revealed nine as the sample size. However, to ensure that the results of each experiment were reliable and replicable, we used more than 20 participants for each experiment and at least 9 participants per subgroup. A total of 98 young healthy participants participated. Seventy-one participants participated in experiment 1, and 27 participants participated in experiment 2. All participants provided written informed consent, and the study was approved by the institutional review board at RIKEN.

### Screening process for eligibility

The screening process for eligibility took place to confirm the following aspects (*87*, *88*). First, the participants were between 18 and 35 years old. Second, on the basis of a self-report questionnaire, anyone who had neurological or psychiatric conditions (e.g., epilepsy, migraine, stroke, chronic pain, major depression, anxiety disorder, and psychotic disorder), was currently using medication, or was suspected of having a sleep disorder (e.g., insomnia, sleep apnea syndrome, hypersomnolence, restless legs syndrome, and parasomnias) was not eligible to participate. Third, participants who had an irregular sleep schedule, i.e., those whose sleep/wake time differed by more than 2 hours between weekdays and weekends, were not eligible. We also ensured that the subjects had not taken a trip to a different time zone during the past six months prior to the experiment. Participants who worked night shifts or who had a habit of taking a nap were not eligible for this study. Fourth, those who failed to complete an MRI safety questionnaire (i.e., claustrophobic, pregnant, or had metallic objects in the body, such as vascular clips, prosthetic valves, metal prostheses, metal fragments, pacemakers, or dental braces) were not eligible for experiment 2. The results of the postconsent questionnaires indicated that none of the participants had sleep problems. Nevertheless, considering that our screening process was based on a self-report questionnaire, we excluded participants whose PSG data showed a deviation from normal, such as sleep-onset REM sleep (see Inclusion and exclusion of data below).

### Inclusion and exclusion of data

Out of 98 participants, 11 participants’ data were excluded from the analyses. Five participants experienced sleep-onset REM periods during their sleep session. Data from another three participants were excluded because they did not reach the threshold for the category task (see Category task below). Three participants’ data were excluded from the analyses because they were mostly awake during the sleep session. The remaining 87 participants data were included for the analyses (n = 61 in experiment 1, n = 26 in experiment 2). In experiment 1, out of 41 participants data that were assigned to the sleep group, considering the differences in brain networks due to handedness (*89*), three participants who were left-handed were excluded from the EEG analyses. Another two participants’ data were removed from EEG analyses due to poor EEG signal quality.

As a result, 36 participants’ data were included in sleep-EEG analyses for stage N2 sleep and twenty three participants were included for slow-wave sleep. In experiment 2, of the 26 participants, MRS data were acquired from the SMA in 22 participants and from the hippocampus in 4 participants. Due to poor shimming in the hippocampus, the quality of the spectral fitting was largely suboptimal. Another dataset taken from the SMA showed poor signal quality. Thus, these five sets of data were excluded from the MRS analyses. One additional dataset contained no usable MRS measurements for the coupling-specific E/I ratio and was therefore removed from the coupling-specific MRS analyses (see **fMRS analysis for sleep microstructures** below). Among 10 datasets that had REM sleep, two datasets were removed from MRS analyses because of no available data for the measurements of the REM-phasic E/I ratios, and the remaining 8 were included for MRS analyses. The final number of datasets included in the MRS analyses for each sleep stage varied depended on how many datasets had valid MRS samples available for that stage (see **fMRS analysis for sleep stages**).

### Experimental design

#### Common procedures

For one week before the experiments started, the participants were instructed to maintain their regular sleep–wake habits until the study was over. The sleep‒wake habits of the participants were monitored by a sleep log and an actigraphy device (MTN-221, Acos, Co., Ltd., Japan; SleepSign Act ver.2, Kissei Comtec, Co., Ltd., Japan). Throughout their participation, the participants were instructed to refrain from taking naps. One day prior to the experiments, alcohol consumption was not allowed. Unusual excessive physical exercise and caffeine consumption were not allowed on the day of the experiments. Approximately one week prior to the main experimental session, an adaptation sleep session was conducted to mitigate the first-night effect (*90–93*). During the adaptation sleep session, the electrodes for the PSG measurement were attached to the participants (see PSG measurement below). Participants went through the same sleep experiments as they did in the main sleep session, without (experiment 1) or with (experiment 2) MRI scans (see MRI measurement below).

#### Experiment 1

The participants were recruited in two phases to equate the number of participants in the NREM + REM and NREM-only groups (*23*, *94*). In the first phase, we employed a passive method in which the duration of the sleep session was fixed to approximately 90 minutes to obtain sleep data passively. Afterward, we classified the participants into the NREM + REM (including REM sleep after NREM sleep) or NREM-only (including only NREM sleep without REM sleep following it) group on the basis of their sleep content. In the second phase, to increase the number of NREM-only group participants, we used a termination method (*23*, *94*). The experimenters scored sleep stages in real time, and the sleep session was terminated when the PSG recording showed a characteristic the ending of an NREM sleep cycle that typically occurred before the onset of REM sleep. These characteristics include sleep stage N1 or N2 reappearing following N3 and EEG becoming desynchronized and decreasing EMG activity after N2 continued for more than 40 minutes without N3 (see PSG measurement below). We continued this method until the number of individuals in the NREM-only group reached at least nine.

Following the adaptation sleep session, the main experimental session took place. The participants arrived at the experimental room at approximately noon. A questionnaire regarding their bedtime and whether they had consumed alcoholic beverages the night before the experiment and the wake-up time of the day of the experiment was given. Afterward, the first half of the task session was performed (see the two-task paradigm below). After the task session, the PSG electrodes were attached, which took approximately 30–45 min (see PSG measurement below). Then, a nap session began in the sound-attenuated, electrically shielded room. After the nap session, there was a 30-min break to ameliorate sleep inertia (Lubin 1976). During the break, a questionnaire was administered to obtain introspection about their sleep during the nap, including subjective sleep-onset latency, subjective wake time after sleep onset, comfort of the environment and occurrence of dreams. After the nap, the second half of the task session, which consisted of a retest of the task, was conducted.

#### Experiment 2

The adaptation and the main experimental sessions for experiment 2 were conducted in a similar manner as in experiment 1, except that magnetic resonance imaging was performed simultaneously with PSG during the sleep session. Shortly after the completion of the first half of the task session, the electrodes for PSG were attached to the participants (see PSG measurement). Afterward, the participants entered the MRI scanner. Anatomical structure measurement, voxel placement and shimming were conducted (see MRI acquisition); then, the MRS measurement started, the room lights were turned off, and the sleep session began. The sleep session lasted for approximately 90 min. After the sleep session, a questionnaire regarding their sleep, the break, and the second half of the task session was administered, as in experiment 1.

The passive method for subject recruitment was performed throughout experiment 2 until the number of participants reached at least 10 for each of the congruent and incongruent groups because the main purpose of experiment 2 was to measure the metabolite concentrations during NREM and REM sleep. Considering the extremely difficult labor in the collection of simultaneous 7-T MRI and PSG data, aiming for the same number of participants for the NREM-REM and NREM-only groups was not feasible.

### Two-task paradigm

The participants performed two tasks: the motor task as a nondeclarative procedural task and the category task as a declarative task. The head and chin of the participants were restrained by a chin rest to precisely maintain a viewing distance of 60 cm. During the main experimental session, the motor task occurred first, followed by the category task before and after an interval. Within each task, the practice session was followed by a postinterval test session. Details for each task are described below.

#### Motor task

The finger-tapping motor-sequence task developed previously (*39*) was used. The experimental design was similar to that in previous studies (*39*, *40*, *49*, *95*). The task requires participants to press four numeric keys on a standard computer keyboard with the fingers of their nondominant hand, repeating a twelve-element sequence as quickly and as accurately as possible. The same sequence [4 – 3 – 2 – 3 – 2 – 2 – 1 – 3 – 1 – 4 – 4 – 1] was used across participants to keep the degree of task difficulty constant because the order of the key presses within the sequence is known to affect the difficulty of movement in this task (*96*). The numeric sequence was always displayed on the screen to exclude any working memory component from the task. There was a practice session prior to the main testing and training session. The sequence for the practice session was [1 – 2 – 3 – 4], because the practice session was conducted for participants to adapt to how to perform the task. The main session consisted of testing and training sessions. The participants first conducted the pretraining test session (pretraining) and then conducted the training session (training). After the training session, they conducted a posttraining test session (posttraining) to investigate the effects of training. After the posttraining test session for the category task, another retest session occurred (data not shown). After an interval, there was a practice session, followed by a postinterval test session (postinterval) to investigate the effects of an interval (wakefulness or sleep). The test sessions consisted of three 30-s blocks with 30-s rest periods between blocks and lasted approximately 3 min. The training sessions consisted of twelve 30-s blocks with 30-s rest periods between blocks and lasted approximately 13 min. We measured the number of correctly typed responses, sequences, and reaction times for each block. Performance improvement by training was defined as the difference between the pretraining performance and the posttraining performance measures, and that by sleep was defined as the difference between the posttraining performance measures and the postinterval performance measures.

#### Category task

In the category task, participants were instructed to memorize the order of the category of each visual stimulus. There were 12 images in four categories, namely, faces, animals, scenes, and objects, in a predetermined order. The order of the categories differed depending on the condition to which the participants were assigned (see <u>Higher-order sequence</u> below; congruent: [object – scene – animal – scene – animal – animal – face – scene – face – object – object – face]; incongruent: [scene – animal – object – object – scene – face – animal – face – object – face – scene – animal]). Participants were presented each image only once, except for the practice session and when training sessions were repeated. Participants were explicitly instructed that the sequence consisted of four categories and that the aim of the task was to memorize the order of the presented sequences. There were two tasks within the category task, the probe and the recall tasks, to facilitate memorization of the sequence. During the probe task, while the images were presented, the participants were instructed to choose an upcoming image from two candidate images. For each block, the probe test appeared 4 times, and participants were presented with a different set of images, with the category sequence remaining the same across all blocks. Participants pressed either 1 or 2 on a keyboard to respond to the probe task. Each time, the answer key was randomly generated to ensure that the category task did not require procedural skills. The interstimulus interval was 0.8 s for the test and training sessions, whereas it was 1.8 s for the practice session. The time limit to respond to the probe task was not set because the aim of the task was to memorize the category order, and responding quickly was not a requisite. The category task consisted of a practice, followed by a pretraining test, training, posttraining, and postinterval test. During the practice and training sessions, the correct image for the probe task was provided after the participant responded, whereas during the test sessions, no feedback was provided. Each test session included 2 blocks. Each training session included at least 6 blocks. The participants repeated the training blocks a maximum of 3 times until they reached an 80% correct rate to ensure that they remembered the sequence. At the end of each test session, participants were instructed to verbally report the sequence of categories. The number of correct responses and the reaction times were measured for the probe task, and the number of correctly reported categories within the sequence was measured for the recall task. The images used for the category task were chosen from websites (e.g., https://unsplash.com, https://pixabay.com, https://www.photo-ac.com/) and modified to match the dimension; the background was also removed. The images in Fig. 1 were selected from Pixabay (https://pixabay.com/photos/) and modified to match the dimension and background as in the actual task setting.

#### Higher-order sequence

To test the effects of higher-order information between the motor and category tasks on behavior and brain activity during sleep, a rule was hidden, inspired by a previous study (*18*). The participants were assigned to either the congruent or the incongruent condition. In the congruent condition, the two tasks shared a higher-order common sequence; that is, each category was assigned to a specific digit in the motor task (1 for Face, 2 for Animal, 3 for Scene, 4 for Object), thus creating a higher-order common sequence. In the congruent condition, there was no such correspondence between the sequences.

### Sleepiness measurement

Before the pretraining and postinterval task sessions, the Stanford Sleepiness Scale (SSS) (*97*) and psychomotor vigilance task (PVT) (*98*) were used to measure sleepiness. The SSS score ranged from 1 (feeling active, vital, alert or wide awake) to 7 (no longer fighting sleep, sleep onset soon or having dream-like thoughts). Participants were asked to choose the scale rating that described their state of sleepiness. In a trial of the PVT, after a fixation screen, a target screen was presented in which a red circle appeared at the center of the screen. The participants were required to press the spacebar on a keyboard as quickly as possible after detection of the circle. The time interval between the fixation screen and the screen with the red circle varied between approximately 1,000 and 10,000 ms. The PVT lasted for approximately 2 min. The reaction time (RT) was log-transformed. The mean RT, the number of lapses, and the lapse threshold were obtained as measurements of behavioral sleepiness (*99*). Lapses were defined as RTs greater than 500 ms. The lapse thresholds at 500 ms were obtained by plotting a cumulative distribution function of RTs across participants for each group. We tested whether each sleepiness measure differed between the congruent and incongruent groups in each experiment.

### MRI acquisition

In experiment 2, MRI data were acquired with a 7-Tesla system (GE SIGNA 7T) using a 64-channel head and neck coil at RIKEN. An anatomical T1-weighted image was acquired using a 3D magnetization-prepared rapid gradient echo (MPRAGE) sequence (TR = 3397.04 ms, TI = 1200 ms, TE = 3.908 ms, FA = 8°, voxel size = 0.7×0.7×0.7 mm^3^, FOV = 320 × 320 × 256 mm). Functional MRS (fMRS) data were acquired using a STEAM sequence (*100*) (TR/TE=2500/13 ms; VOI=30 × 30 x 8 mm^3^). A CHESS (*101*) was used to simultaneously suppress the water signal. MRS data were acquired from the right motor area, covering the trained SMA that was contralateral to the left hand used for the motor task (*49*). An fMRS run had 240 volumes of metabolite signal and 16 volumes of unsuppressed water signal, taking approximately 10 min to acquire. The unsuppressed water signals were later averaged and used as a standard water concentration reference (*102–104*). The same run was repeated 9 times if possible, resulting in 9 runs in total, during the nap session. When the TR was 2.5 s, owing to phase cycling = 2, the minimum practical temporal resolution was 5 s. Cardiac and respiratory cycles were simultaneously recorded using a fingertip PPG sensor and respiratory belt (Biopac, US) with a 100-Hz sampling rate. MRI data were acquired simultaneously with PSG data (see the PSG measurement section below). Cushions and gauze were used to stabilize the subjects’ heads to reduce discomfort and head motion. Back and knee cushions were used upon the participants’ request to further reduce discomfort. Several blankets were used to keep the participants warm and to initiate sleep during the scan (*88*).

### PSG measurement

#### Experiment 1

PSG data were acquired using a BrainAmp Standard or actiChamp EEG amplifier (Brain Products, Gilching, Germany) with a sampling rate of 500 Hz and 64-channel EEG cap, except for 4 participants’ data, where a 32-channel EEG cap was used (EasyCap, Wörthsee, Germany). For all the main sessions, the BrainAmp Standard amplifier was used. The hardware bandpass filters were set to a range of 0.016–250 Hz, with a roll-off of 30 dB/octave at high frequency. The electrode layout followed the extended international 10-20 system. Electrooculograms (EOGs) were obtained from the outer canthi of both eyes to measure horizontal eye movements and above and below the right eye for vertical eye movements using bipolar electrodes. Electromyograms (EMGs) were measured from two bipolar electrodes placed on the chin. To record the electrocardiogram (ECG), two bipolar electrodes were placed, one on the clavicle and the other on the rib bone. For 16 participants’ data, Fz was used as the reference electrode and Fpz was used as the ground electrode, whereas for 33 participants’ data, FCz was used as the reference electrode, whereas Fpz was used as the ground electrode. EEG data were transferred outside the scanner room through fiber optic cables to a computer running the BrainVision Recorder software (Brain Products, Gilching, Germany). We used active electrodes for EEG. The impedance of all EEG electrodes was approximately 15 kΩ. The active electrodes included an integrated impedance converter, which allowed them to transmit the EEG signal with significantly lower levels of noise than traditional passive electrode systems do. Passive electrodes were used for the EOG, EMG, and ECG measurements. The impedance level was approximately 10 kΩ.

#### Experiment 2

PSG data were acquired using a BrainAmp MRplus and BrainAmp ExG MR amplifier (Brain Products, Gilching, Germany) with a 5-kHz sampling rate and a custom-made MR-compatible 32-channel EEG cap for 7-Tesla MRI (BrainCap MR7flex 32channnel Cap, Brain Products, Gilching, Germany). The hardware bandpass filters were set to a range of 0.016–250 Hz, with a roll-off of 30 dB/octave at high frequency. The electrode layout followed the extended international 10-20 system with additional drop-down channels for recording the ECG, EOG, and chin EMG. EEG was measured from 29 electrodes. Two EOG electrodes were placed at the outer canthi. EMG was measured bipolarly from the mentum. ECG signals were measured from an electrode placed at a lower shoulder blade. Four carbon wire loops were attached to the cap. FCz was used as the reference electrode, whereas AFz was used as the ground electrode. EEG data were transferred outside the scanner room through fiber optic cables to a computer running the BrainVision Recorder software (Brain Products, Gilching, Germany). MR-EEG scanner clocks were synchronized (Brain Products Synchbox) for all PSG data acquisitions, and the onset of every scanner repetition time (TR) period was recorded in the EEG data for MR gradient artifact correction. The impedance of all electrodes was initially maintained below 10 kΩ and was maintained below a maximum of 20 kΩ for the duration of the study.

### PSG data preprocessing

In experiment 2, PSG data were acquired simultaneously with MRI data continuously for approximately 90 minutes. The PSG data were contaminated by two types of artifacts, MRI-related gradients and ballistocardiogram (BCG) artifacts. To remove the MRI-related gradient artifacts from the PSG data caused by the switching of the strong magnetic field gradient, the PSG signals were first corrected using BrainVision Analyzer2 (Version 2.2.0, Brain Products GmbH, Gilching, Germany) with the standard average artifact subtraction (AAS) method. The AAS method uses sliding window templates formed from the averages of 21 TRs, which are subtracted from each occurrence of the respective artifacts for each electrode (*88*, *105*, *106*). The gradient-artifact corrected data were subsequently bandpass filtered (EEG: 0.1–50 Hz with a notch filter of 50 Hz) and downsampled to 250 Hz. Following the gradient artifact correction, cardiac R-peaks were automatically detected from the ECG recording in BrainVision Analyzer2, checked visually and corrected manually when necessary. These R-peak events were used to inform BCG correction. The artifacts were further corrected via the AAS method with sliding window templates formed from the averages of 21 R-peaks in BrainVision Analyzer2 (*88*, *107*, *108*). Independent component analysis (ICA) was applied to remove residual noise. These procedures were conducted for the entire duration of the fMRS scan to obtain artifact-removed PSG data for further processing (see Sleep stage scoring below).

### Sleep stage scoring

Sleep staging was conducted on the artifact-removed PSG data following the standard criteria (*109*, *110*). Prior to sleep staging, PSG data were further bandpass filtered between 0.3 and 40 Hz for EEG, EOG and ECG and between 10 and 50 Hz for EMG to remove low-frequency drift and high-frequency noise and were rereferenced to the linked mastoids (TP9 & TP10). The sleep stages were scored every 30 s in conjunction by trained sleep scorers (XL, AS, and MT). We obtained sleep parameters, including the total sleep time and duration of wakefulness (W), NREM 1 (N1), NREM 2 (N2), NREM 3 (N3), and REM (REM) (table S1). Epochs with motion artifacts and arousal events (see below) were marked and removed from further analyses (*23*, *35*, *88*). An arousal event was defined as an abrupt shift in EEG frequency, which may include theta, alpha and/or frequencies greater than 16 Hz (but not spindles) that were at least 3 seconds in duration, with at least 10 seconds of stable sleep preceding the change, in accordance with the standard AASM criteria (*110*). Arousal events were manually scored by visually checking the PSG signals. The consensus of the arousal scoring was used, and any 30-s epochs containing arousal events were removed from further analyses. Motion artifacts were defined as an epoch containing artifacts that lasted longer than 1 s or a participant showing motion (experiment 1 and 2) in the video recording (experiment 2).

### EEG source reconstruction

In experiment 1, to estimate the EEG activity in each brain region of interest (see ROIs below), a scalar linearly constrained minimum variance (LCMV) beamforming analysis (*63*, *111–113*) was applied to the EEG data. All EEG source reconstruction processes were performed in Brainstorm software (*114*). The standard template of EEG electrode positions was coregistered with the template ICBM152 anatomical image (*115*, *116*). The forward model was computed using OpenMEEG software implemented in Brainstorm (*117*). A 3-layer boundary element method (BEM) head model consisting of brain, skull, and scalp surfaces with conductivity values of 0.33, 0.0165, and 0.33 S/m, respectively (*118*), was used to reconstruct broadband (1–100 Hz) time courses of neural activity in Brainstorm (*114*). The EEG source signal of each vertex location on the cortical surface was reconstructed. The source-reconstructed data were used for sleep spindle detection (see Sleep event detection below).

### Regions of interest (ROIs) for experiment 1

In experiment 1, we performed source reconstruction for the following predetermined four regions of interest (ROIs) as potential waypoints for memory transfer, including the right supplementary motor area (SMA), the parietal cortex, the medial prefrontal cortex (mPFC), and the hippocampus (fig. S1), on the basis of earlier studies (*13*, *48–55*). The right SMA, which was contralateral to the trained hand, is a core motor region involved in finger-tapping tasks (*48–50*) and is composed of a part of the precentral sulcus and the superior frontal gyrus (*51*, *52*). The parietal cortex, which is involved in movement chunking and sequence representations, was defined as the superior parietal gyrus (*53*). The hippocampus was included as a hub for episodic memory processing and the extraction of novel regularities (*119*). The media prefrontal cortex, which is involved in abstract rule representations and the redistribution of memories (*54–56*), comprised the anterior cingulate and superior frontal gyrus-medial regions (*55*, *56*). These ROIs were determined anatomically according to the automated anatomical labeling atlas 3 (*120*).

### Sleep event detection

#### Slow waves

Slow waves were detected semiautomatically on the scalp EEG signals as follows. First, the scalp EEG signals were bandpass filtered between 0.1 and 4 Hz with a zero-phase 4th-order Butterworth filter. Then, slow waves were detected automatically (*121*) using the Wonambi software (https://wonambi-python.github.io/) from the AF4, F2, F4, FC2, FC4, and C4 electrodes during the N2 and N3 stages. Only the waves that met the following three criteria were selected for the detection of well-defined, large-amplitude slow waves: 1) a positive-to-negative zero crossing and a subsequent negative-to-positive zero crossing separated between 0.3 s and 1 s, 2) a negative peak between the two zero crossings with voltages smaller than −80 μV, and 3) a peak-to-peak amplitude greater than 140 μV. The automatically detected slow waves were then reviewed and removed by an experimenter if necessary. The number of slow waves was measured for each electrode, and the electrode that had the maximum number of slow waves was used as the most sensitive electrode for slow wave detection. The density of slow waves was measured as the total number of slow waves divided by the duration of N2 or N3.

#### Sleep spindles

Sleep spindles were semiautomatically detected (*122*) using Wonambi software (https://wonambi-python.github.io/) from source-space ROIs (experiment 1) or FC4, which is near the SMA (*123*) (experiment 2), during the N2 and N3 stages as follows. In short, a bandpass filter was applied between 0.3 and 40 Hz. The root mean square (RMS) of the filtered signals was subsequently calculated at every sample point using a moving window of 0.2 s. The threshold for spindle detection in the RMS signal was set to 1.5 standard deviations of the filtered signal for each electrode. A spindle was detected when the RMS signal remained above the threshold for 0.5–3 s, after which the beginning and end of the spindle were marked at the threshold crossing points. The automatically detected spindles were reviewed and removed by an experimenter if necessary. The density of sleep spindles was measured as the total number of sleep spindles divided by the duration of N2 or N3.

#### Slow wave–sleep spindle coupling events

We classified a ‘coupled’ slow-wave event if the onset of a sleep spindle was detected within 750 ms from the onset of the negative peak of a slow wave (*124*, *125*). Otherwise, the slow wave was classified as an uncoupled slow wave. The density of slow wave‒spindle coupling was measured as the total number of events divided by the duration of N2 or N3. In experiment 2, to test the neurochemical changes associated with coupling, the N2 and N3 sleep stages were further classified into coupled or uncoupled periods every 5 seconds. If a 5-second period included a slow wave–spindle coupling event, the period was classified as a coupled period. If no coupling event was detected, the period was classified as an uncoupled period. Because of the small amount of slow-wave sleep in experiment 2, we combined N2 and slow-wave sleep and measured the mean value.

#### REM phasic and tonic periods

Rapid eye movement events were semiautomatically detected from two EOG signals (*88*, *126*) during REM sleep as follows. To separate rapid eye movements from slow eye movements (SEMs), two channels of EOG data are filtered using a fourth-order Butterworth bandpass filter with cutoffs at 1 and 9 Hz to yield the filtered EOG data. The candidate REM sleep events were generated as the negative instantaneous product of the two filtered EOG sequences, and the REM sleep events were detected on the basis of their amplitude threshold. The amplitude threshold was arbitrarily determined on the basis of each individual dataset, since owing to differences in the setup and/or the quality of the artifact removal, there were individual differences in the amplitude of the EOG data. In experiment 2, these were detected every 5-second epoch to match the temporal resolution of the MRS data (see fMRS analysis for sleep microstructures below). Epochs containing at least two rapid eye movements were classified as a phasic period, and epochs having fewer than two rapid eye movements (one or none) were classified as a tonic period.

### Density measurement

For sleep microstructures (slow waves, sleep spindles, coupled slow wave–spindles, REM phasic, REM tonic periods), the number of events or periods per minute was measured as the density.

### Coherence analysis

#### NREM sleep

For coherence analysis during NREM sleep, five source signals of ROIs (right SMA, the parietal cortex, mPFC, and the left and right hippocampus) time-locked to coupled and uncoupled events (see <u>Slow wave–sleep spindle coupling events</u> in Sleep event detection above) were used. The coherence values were calculated between 0 and 1 s from 5 pairs and averaged in the frequency band between 0.5 and 2 Hz corresponding to the slow-wave frequency for each pair of each coupling event type using the Fieldtrip toolbox (Fieldtrip-20191014; http://www.ru.nl/neuroimaging/fieldtrip) (*127*). The coherence value was Fisher *z* transformed before it was averaged across participants in each group for each event. Slow-wave spindle-specific coherence values were measured by determining the difference between the coherence values for coupled and events.

#### REM sleep

For coherence analysis during REM sleep, first, every 3-s epoch during REM sleep was classified into a phasic or tonic period (see <u>REM phasic and tonic periods</u> above). The coherence values from 5 pairs were calculated for every 3-s epoch and averaged in the frequency band between 5 and 9 Hz, corresponding to the theta band for each pair of each coupling event type using the Fieldtrip toolbox (Fieldtrip-20191014; http://www.ru.nl/neuroimaging/fieldtrip) (*127*). The coherence values were Fisher z transformed and then averaged across participants in each group for each phasic and tonic period. REM-specific coherence values were measured by taking the difference between the coherence values for the phasic and tonic periods.

### fMRS analysis for sleep microstructures

MRS data were analyzed using FSL-MRS (https://open.oxcin.ox.ac.uk/pages/fsl/fsl_mrs/) (*128*). For each 10-min MRS run, the metabolite signals were segmented into 5-second epochs (120 epochs per run). For each 5-second epoch, after the FIDs were averaged and the coils were combined, eddy current correction, water signal removal, and phase correction were performed before fitting. For the water reference signals, filtering and phase correction were applied. Tissue segmentation was conducted using individual T1 data to produce water-scaled molarity and molality concentrations. For fitting, the LCModel (*129*) default basis set (TE=15 ms) was used, and the water reference data that were taken from the same run were applied for water scaling for each of the epochs. We measured the GABA and glutamate concentrations (mol/kg) (*130*) and E/I ratios by taking the glutamate-to-GABA ratios for each 5-second epoch.

Because we used a shorter MRS segment to extract spectra than in previous studies (*19*, *23*), we adopted a relatively conservative approach. First, MRS data quality was examined for each 5-second epoch (see MRS data quality below), and only the segments with a Cramér‒Rao lower bound smaller than 20% for both glutamate and GABA were included. Second, any epochs containing motion artifacts were excluded (*23*, *88*). Third, any epochs including arousals were excluded (*23*, *88*). The remaining segments were included for further analyses.

The epochs corresponding to the N2 and N3 sleep stages were further classified into slow wave–spindle coupled or uncoupled periods (see <u>Slow wave–sleep spindle coupling</u> above), and the GABA and glutamate concentrations and E/I ratios were averaged for each of the coupled and uncoupled periods. For REM sleep, the levels of GABA and glutamate and the E/I ratio were averaged for each phasic and tonic period.

### fMRS analysis for sleep stages

To coregister between sleep stages and MRS measurements, the metabolite signals were segmented into 30-second epochs (20 epochs per run). All the preprocessing steps were the same as those for the sleep microstructures. Only the segments with a Cramér‒Rao lower bound smaller than 20% for both glutamate and GABA were included. Any epochs containing motion artifacts or arousals were excluded (*23*, *88*). The remaining segments were included for further analyses. We measured the GABA and glutamate concentrations (mol/kg) and E/I ratios by taking the glutamate-to-GABA ratios for each 30-second epoch to match the temporal resolution of the sleep stage scoring. The concentrations of GABA and glutamate and the E/I ratio were averaged for each sleep stage.

### MRS data quality

We conducted three types of MRS quality checks, namely, water linewidth (Hz), NAA linewidth (Hz), and Cramer–Rao lower bounds, as shown in table S6. We compared these values between the consistent and inconsistent groups to test whether the MRS data quality was comparable between the groups. The results indicated that there was no systematic bias in the MRS data quality. Thus, the MRS data in these groups were sufficiently reliable for a fair comparison.

#### Water linewidth

The linewidth for water was obtained during the prescan of MRS. The T2* of the water in the voxel was determined by fitting the envelope of the free induction decay with a single exponential decay. Then, T2* was converted into the linewidth (*131*).

#### NAA linewidth

The linewidth for the NAA, which determines the resolution available to discern spectral features (therefore, the lower the better), was noted for each MRS run for each participant. We confirmed that the NAA linewidth was less than 25 Hz for all participants.

#### Cramer–Rao lower bounds

The Cramer–Rao lower bounds (or %CRLB) were used as a measure of fitting error (therefore, the lower the better). A commonly accepted Cramer–Rao lower bound criterion of 20% was chosen to reject low-quality signals (*132*). The mean %CRLB values for Glx and GABA were similar to those reported in previous studies (*19*, *23*).

### Correlation analysis between the E/I ratio and delta power

To compute delta power every 30 s, time-frequency analysis using a multitaper approach (*62*, *63*) was applied to the whole EEG signal from the right SMA using the Fieldtrip toolbox (*127*) (http://www.ru.nl/neuroimaging/fieldtrip). Time windows of 0.8 s in duration were moved across the data in steps of 50 ms, resulting in a frequency resolution of 1.25 Hz, and the use of three tapers resulted in a spectral smoothing of ±2.5 Hz. The calculated time–frequency representations (TFRs) were then transformed in the scale of dB by 10*log10. Then, the time–frequency power less than 4 Hz (delta band frequency) for every 30 s was averaged. Pearson’s correlation coefficients between the time course of delta power fluctuations and E/I ratios were subsequently measured.

### Statistical tests

The alpha level was set as 0.05. All tests conducted were two-tailed. The Shapiro‒Wilk test was used to test the normality of the data. When the data were regarded as normally distributed, Analysis of Variance (ANOVA), two-tailed independent-samples or one-sample t tests were conducted. When the data were regarded as nonnormally distributed, the Kruskal‒Wallis test, Mann–Whitney U test or Wilcoxon signed-rank test was used. To control for multiple comparison problems, false discovery rate (FDR) corrections using the Benjamini and Hochberg (1995) method (*133*) were applied for behavioral and coherence analyses. We included a 95% CI for significant *t* tests. Statistical tests were performed in MATLAB (R2023a and R2025b, MathWorks, Inc.).

## Acknowledgements

The authors thank B.D. Fulcher for feedback on an earlier draft, A. Saotome, M. Yamaji, and M. Ogata for support during data collection, M. Matsuoka, A. Kitabori, T. Miura and S. Shoji for administrative supports, A. Ito-Ishida for discussion on the project, Y Miyashita-Fujisawa for support during pilot testing, T. Warbrick and C. Jäger for technical advice on EEG setup in 7 Tesla MRI, S. Someya, T. Uehara, and A. Minohata for help with EEG system. We used generative AI to check grammatical errors and typos in some parts.

## Author Contributions

MT designed the research. XL, MU, RK, CS, KU, MF, RAW, and MT performed the experiments. XL, MU, SA, RAW, and MT analyzed the data. XL, MU, RAW and MT wrote the paper. All confirmed the content.

## Data, code, and materials availability

Source data will be provided with the paper. The original data created for the study will be available in the RIKEN CBS Data Sharing Platform (https://neurodata.riken.jp/). No custom algorithm or software was used to that is central to the present findings. The codes and scripts that were used to generate the results of this study are available from the corresponding author upon request.

## Supplementary Materials

**Fig. S1.**
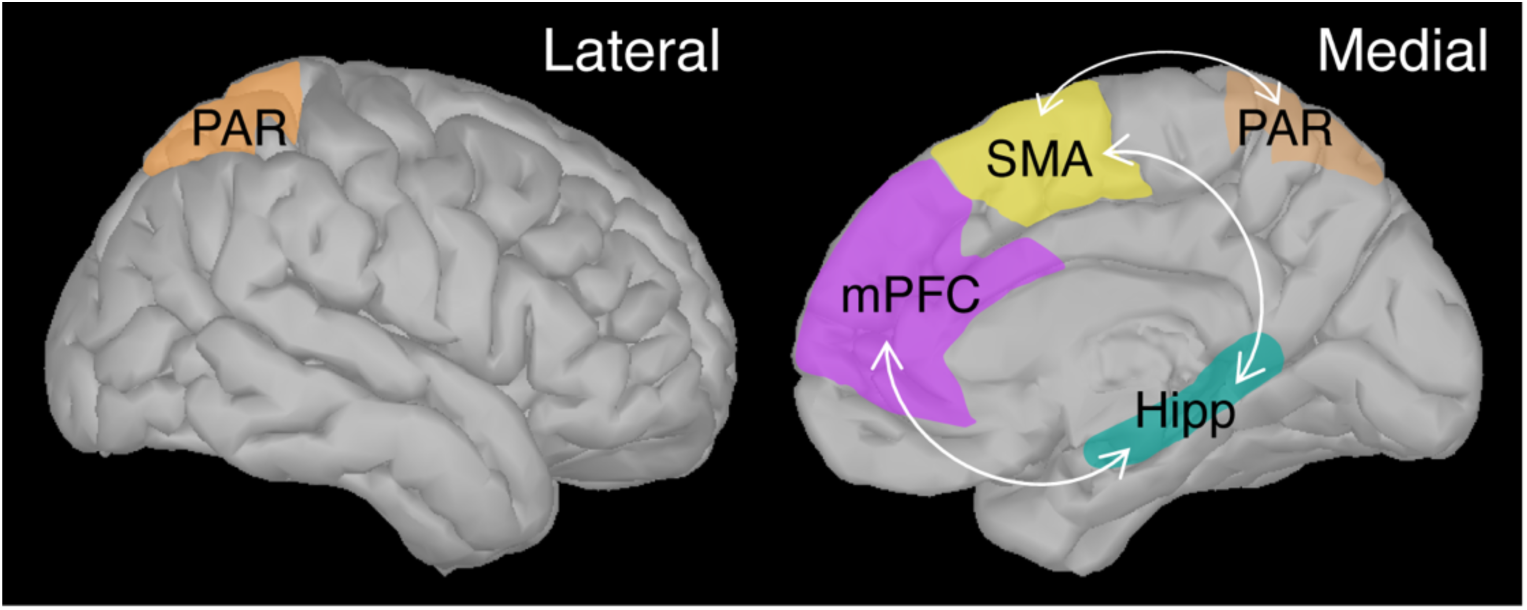
Regions of interest. Lateral (left) and medial (right) images of the right hemisphere. We selected four regions of interest (ROIs) for memory transfer, including the SMA (yellow), the superior parietal cortex (orange), the hippocampus (green), and the mPFC (pink), on the basis of earlier studies (*13*, *48–55*). The right SMA, which was contralateral to the trained hand, is a core motor region involved in finger-tapping tasks (*48–50*) and is composed of a part of the precentral sulcus and the superior frontal gyrus (*51*, *52*). The parietal cortex, which is involved in movement chunking and sequence representations, was defined as the superior parietal gyrus (*53*). The hippocampus was included as a hub for episodic memory processing and the extraction of novel regularities (*119*). The media prefrontal cortex, which is involved in abstract rule representations and the redistribution of memories (*54–56*), comprised the anterior cingulate and superior frontal gyrus-medial regions (*55*, *56*). These ROIs were determined anatomically according to the automated anatomical labeling atlas 3 (*120*). Among the possible pairs, five pairs (white arrows), i.e. mPFC–hippocampus (left and right) (*54–56*), right SMA–parietal (*53*), right SMA–hippocampus (left and right) (*48–50*) were included for coherence measurement based on prior studies (see Regions of Interests in Materials and Methods). The left and right hemispheres for the mPFC and parietal regions are averaged.

**Fig. S2.**
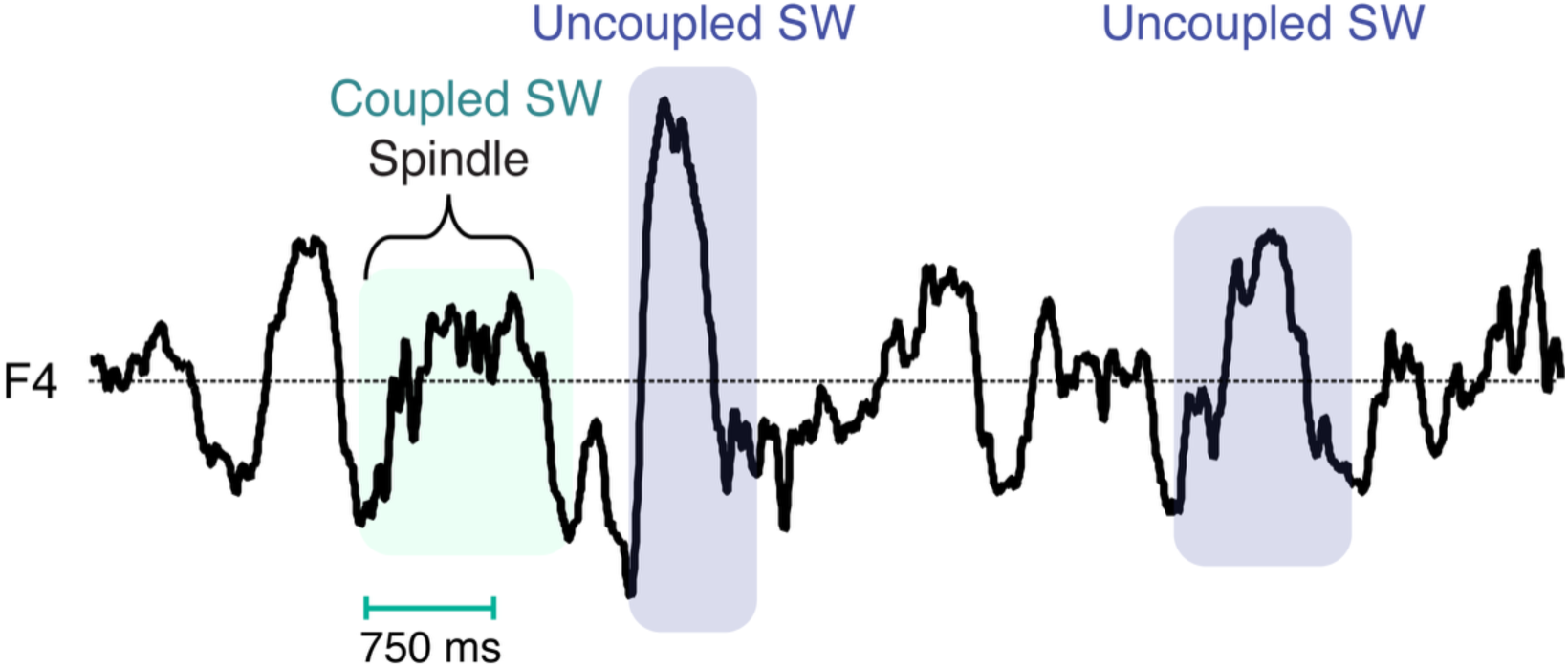
Slow-wave spindle coupled and uncoupled events. We classified a ‘coupled’ slow-wave event if the onset of a sleep spindle was detected within 750 ms from the onset of the negative peak of a slow wave (*124*, *125*). Otherwise, the slow wave was classified as an uncoupled slow wave.

**Fig. S3.**
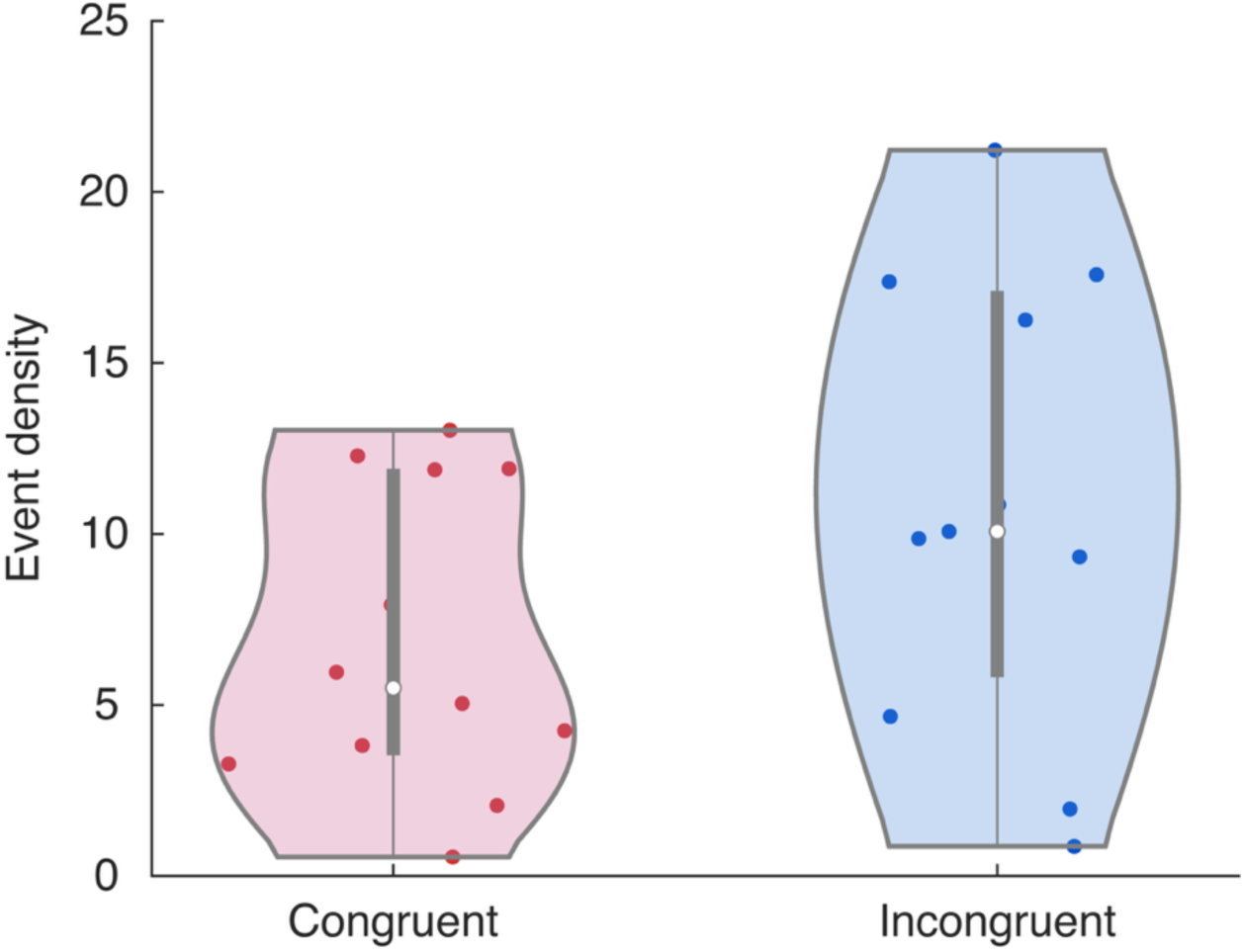
Slow wave density. There was no signficant difference between the conditions (two-tailed independent-samples *t* test, Congruent *n* = 12, Incongruent *n* = 11, *t_21_* = −1.7432, *p* = 0.0959). The number of events per minute was measured as the density.

**Fig. S4.**
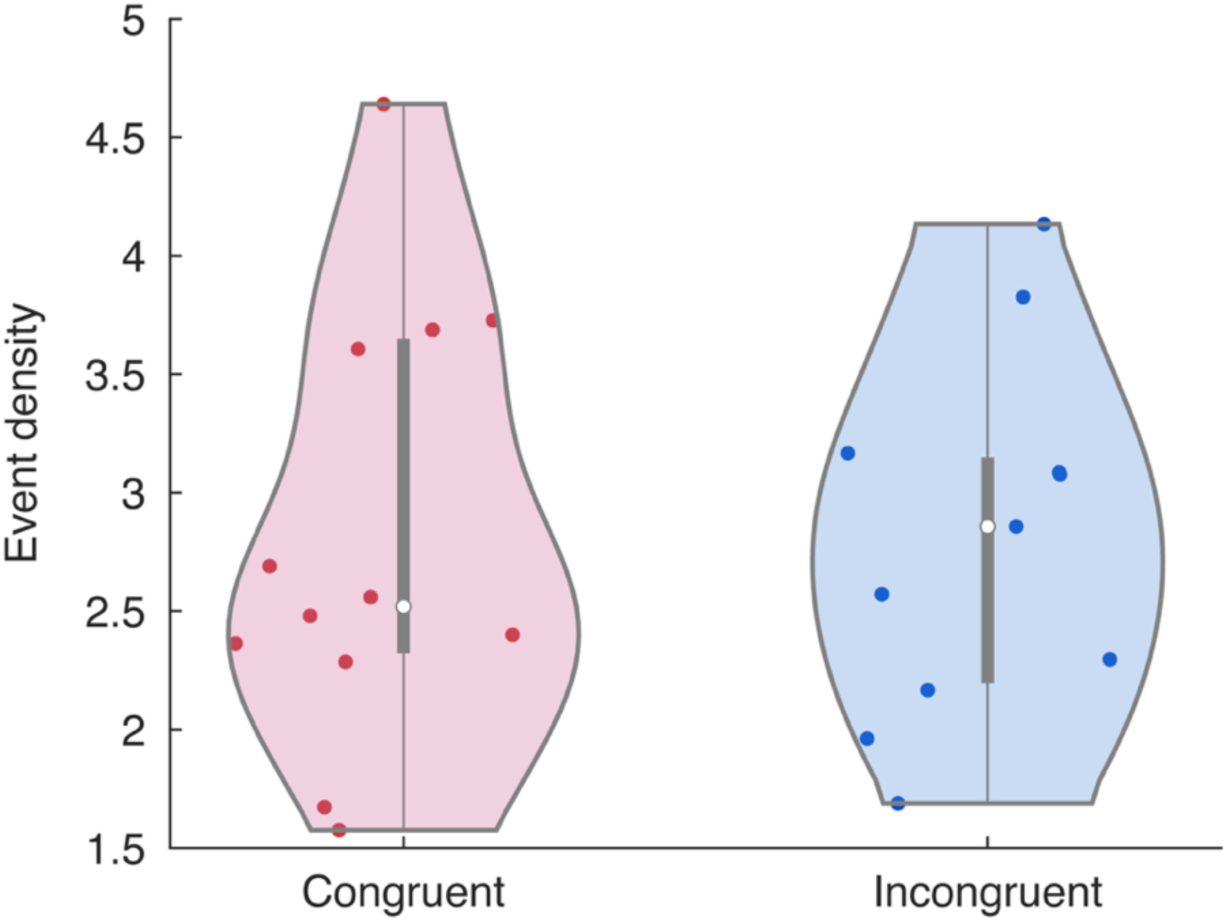
Spindle density. There was no signficant difference between the conditions in the spindle desntiy detected from the right SMA (two-tailed independent-samples *t* test, Congruent *n* = 12, Incongruent *n* = 11, *t_21_* = 0.0127, *p* = 0.9900). The number of events per minute was measured as the density.

**Fig. S5.**
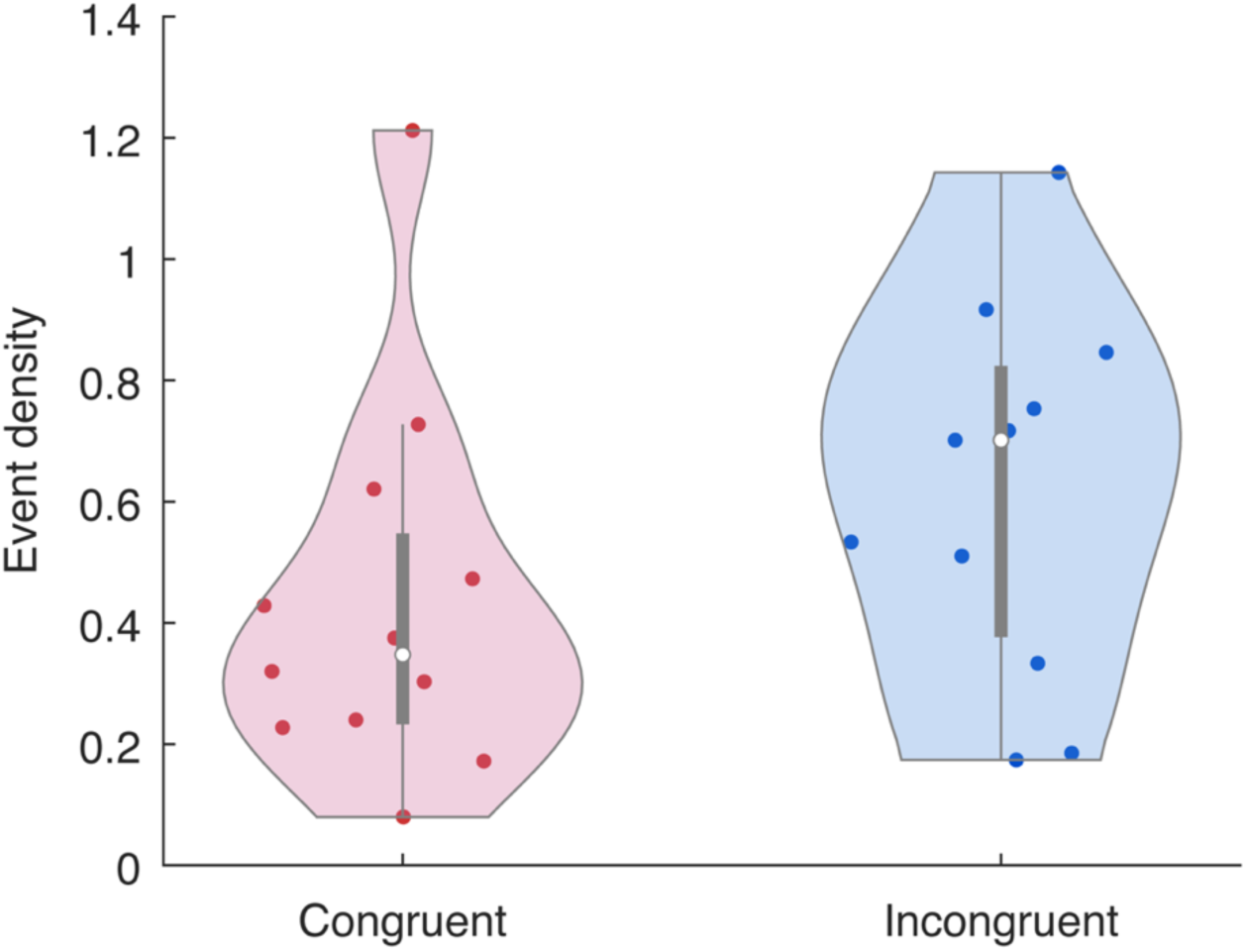
Slow wave-spindle coupling density. There was no significant difference between the conditions in the slow-wave spindle coupling density detected from the right SMA (two-tailed independent-samples *t* test, Congruent *n* = 12, Incongruent *n* = 11, *t*21= −1.4684, *p* = 0.1568). The number of events per minute was measured as the density.

**Fig. S6.**
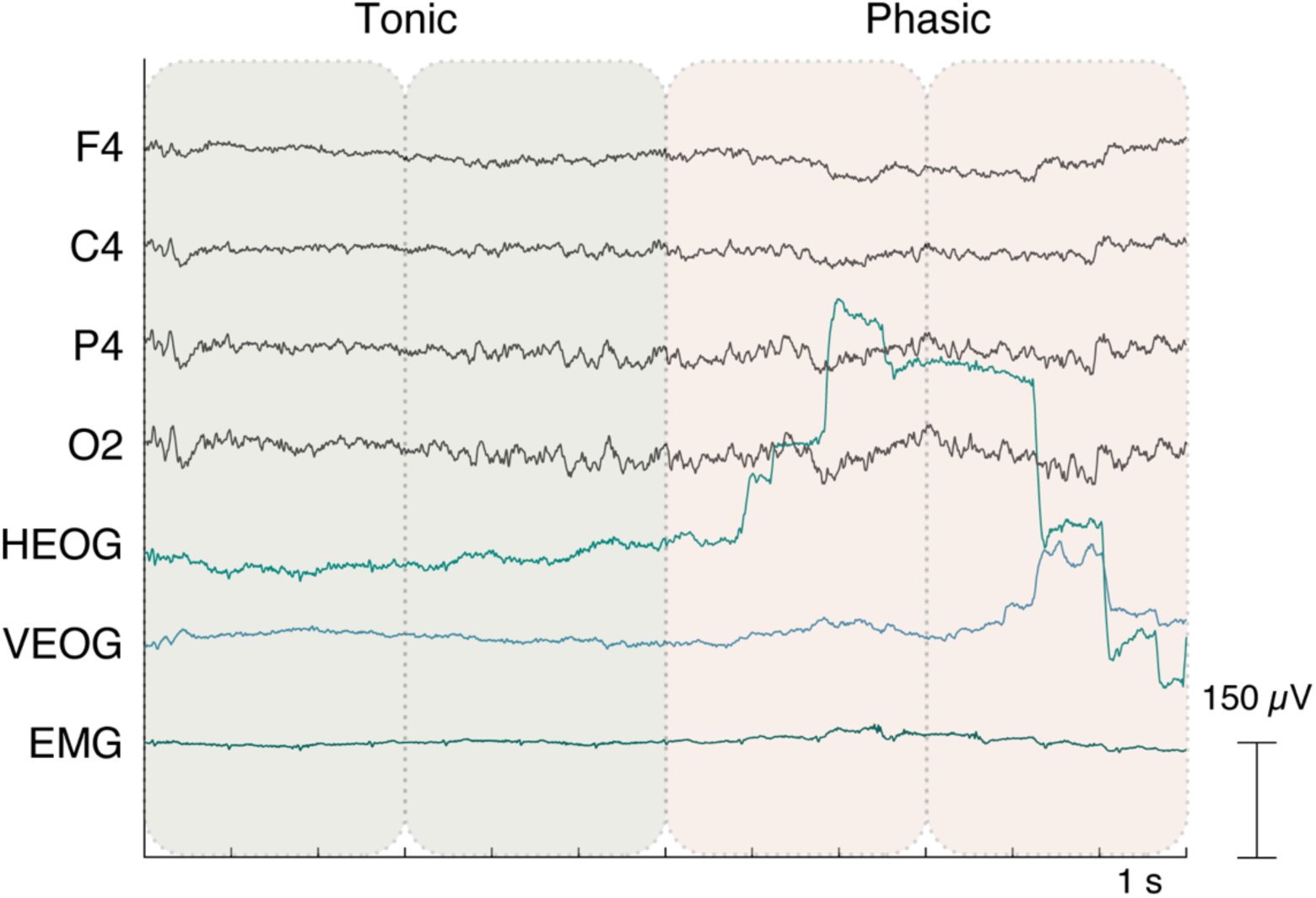
REM-phasic and tonic periods. Rapid eye movement events were semiautomatically detected from two EOG signals (*88*, *126*) during REM sleep as follows. Epochs containing at least two rapid eye movements were classified as a phasic period, and epochs having fewer than two rapid eye movements (one or none) were classified as a tonic period.

**Fig. S7.**
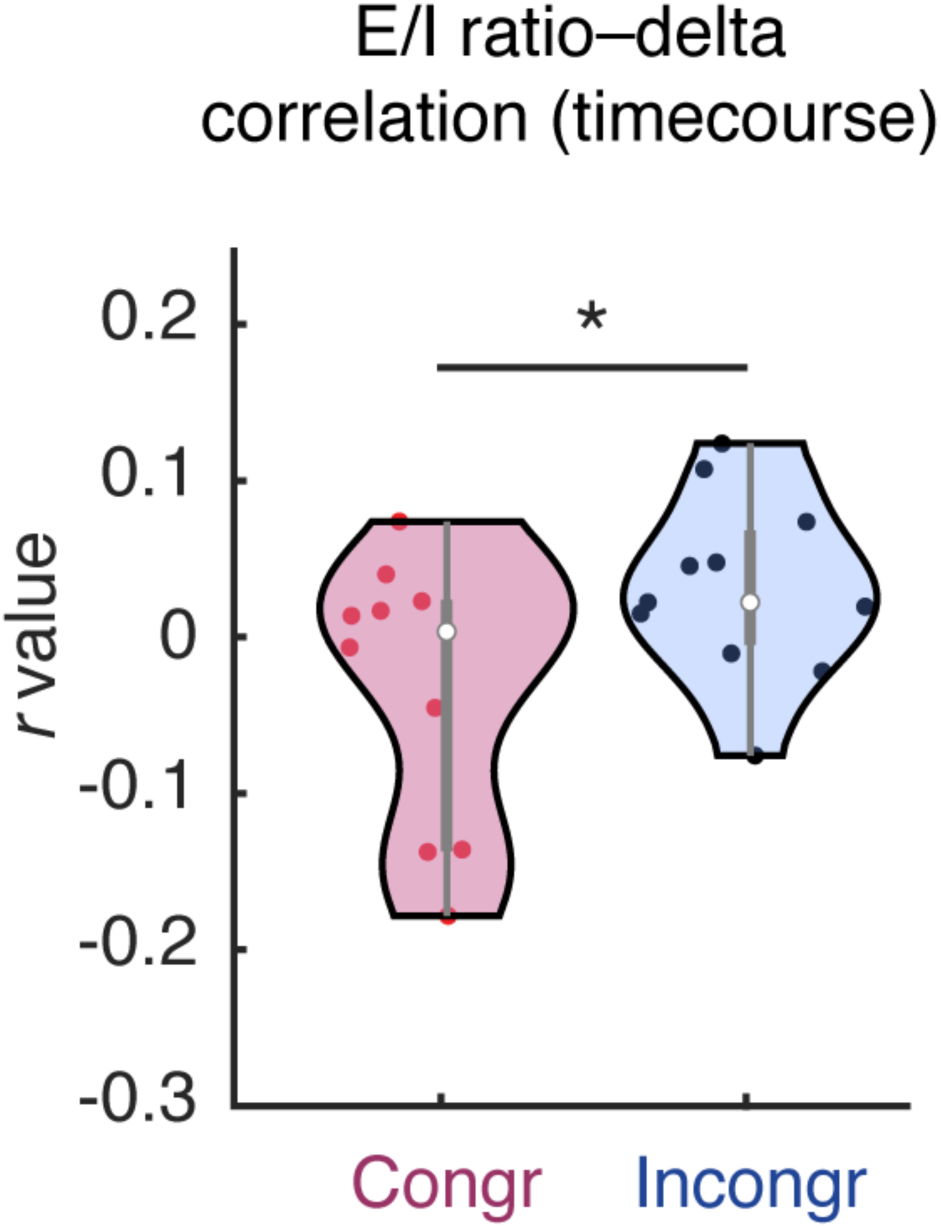
Correlation coefficients between time course delta power and E/I ratio changes. Multitaper analysis was performed on EEG data from the right SMA, where the E/I ratio was measured every 30 s. Pearson’s correlation coefficients between the time course of delta power fluctuations including all the sleep stages and E/I ratios were measured. There was a significant difference in the correlation coefficient between the congruent and the incongruent conditions (two-tailed independent samples *t*-test, *n* = 21, *t*_19_ = −2.2945, *p* = 0.0333, 95% *CI* = [-0.1895, - 0.0087]). The congruent condition shows a more negative correlation coefficient than the incongruent condition.

**Table S1.**
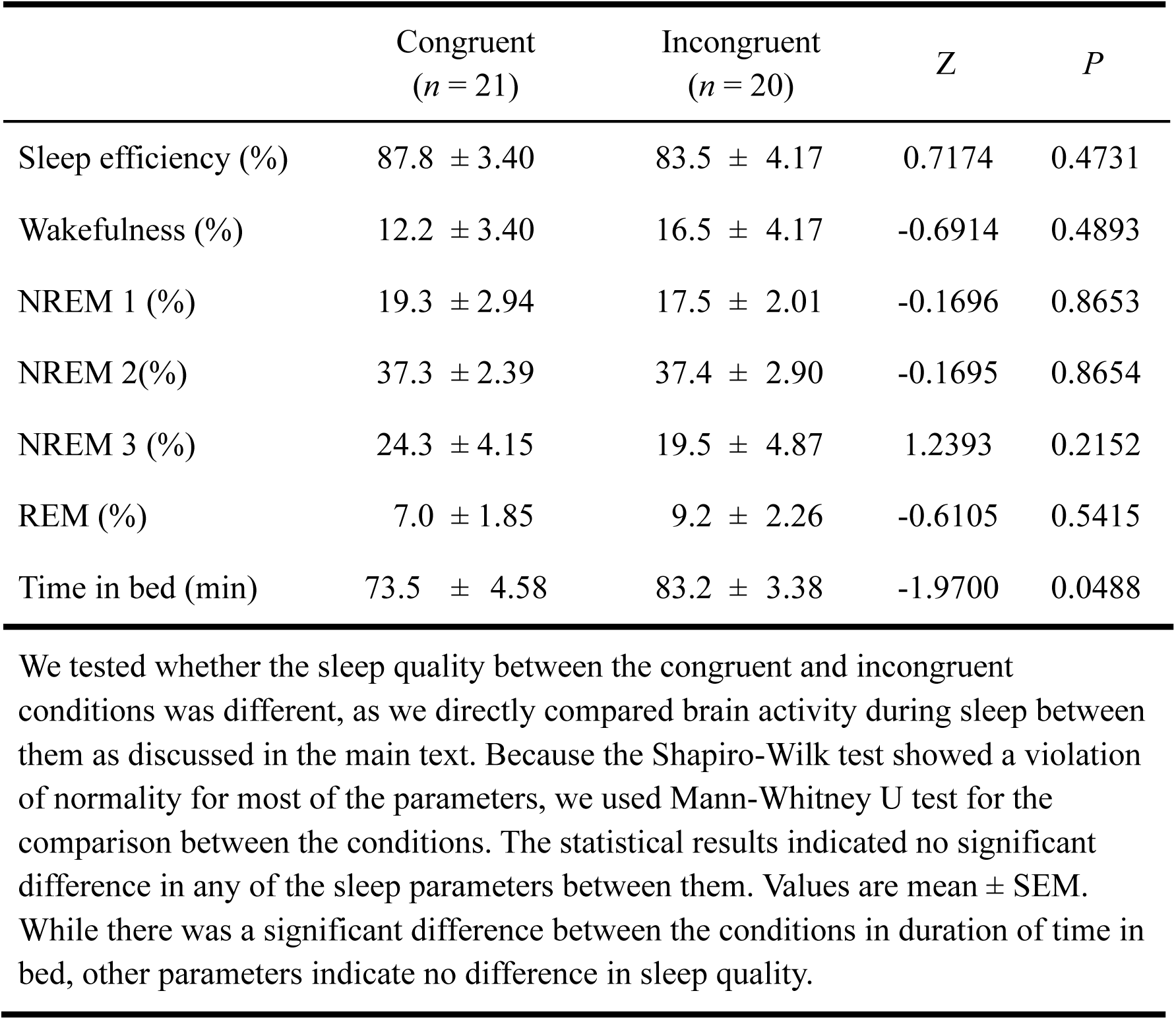
Sleep parameters for experiment 1.

|  | Congruent<br>(*n* = 21) | Incongruent<br>(*n* = 20) | *Z* | *P* |
| --- | --- | --- | --- | --- |
| Sleep efficiency (%) | 87.8 ± 3.40 | 83.5 ± 4.17 | 0.7174 | 0.4731 |
| Wakefulness (%) | 12.2 ± 3.40 | 16.5 ± 4.17 | -0.6914 | 0.4893 |
| NREM 1 (%) | 19.3 ± 2.94 | 17.5 ± 2.01 | -0.1696 | 0.8653 |
| NREM 2(%) | 37.3 ± 2.39 | 37.4 ± 2.90 | -0.1695 | 0.8654 |
| NREM 3 (%) | 24.3 ± 4.15 | 19.5 ± 4.87 | 1.2393 | 0.2152 |
| REM (%) | 7.0 ± 1.85 | 9.2 ± 2.26 | -0.6105 | 0.5415 |
| Time in bed (min) | 73.5 ± 4.58 | 83.2 ± 3.38 | -1.9700 | 0.0488 |

We tested whether the sleep quality between the congruent and incongruent conditions was different, as we directly compared brain activity during sleep between them as discussed in the main text. Because the Shapiro-Wilk test showed a violation of normality for most of the parameters, we used Mann-Whitney U test for the comparison between the conditions. The statistical results indicated no significant difference in any of the sleep parameters between them. Values are mean ± SEM. While there was a significant difference between the conditions in duration of time in bed, other parameters indicate no difference in sleep quality.

**Table S2.** Category task performance for experiment 1.

| Condition | Measure | Pre-training | Post-training | Post-interval |
| --- | --- | --- | --- | --- |
| Sleep Congruent<br>(*n* = 21) | Serial recall | 10 (8-12) | 12 (12-12) | 12 (12-12) |
|  | Probe task (RT) | 0.37±0.17 | -0.64±0.12 | -0.59±0.11 |
|  | Probe task (resp) | 6 (5-8) | 8 (8-8) | 8 (7-8) |
| Sleep Incongruent<br>(*n* = 22) | Serial recall | 8 (5-10) | 12 (12-12) | 12 (12-12) |
|  | Probe task (RT) | 0.11±0.2 | -0.46±0.14 | -0.62±0.16 |
|  | Probe task (resp) | 6 (5-6) | 8 (7-8) | 8 (7-8) |
| Wake Congruent<br>(*n* = 10) | Serial recall | 10 (5-12) | 12 (12-12) | 12 (12-12) |
|  | Probe task (RT) | 0.34±0.2 | -0.73±0.28 | -0.55±0.18 |
|  | Probe task (resp) | 5 (3-8) | 7.5 (5-8) | 8 (7-8) |
| Wake Incongruent<br>(*n* = 10) | Serial recall | 8.5 (5-12) | 12 (12-12) | 12 (10-12) |
|  | Probe task (RT) | -0.04±0.21 | -0.69±0.09 | -0.29±0.28 |
|  | Probe task (resp) | 6 (3-8) | 8 (7-8) | 7.5 (7-8) |

Serial recall, the number of correctly reported category in the serial recall test (median and 95% CI). RT, reaction time (log, mean ± SE). Resp, the number of correct responses (median and 95% CI). Although the statistical results indicated a significant difference among the groups in the serial recall task (the Kruskal–Wallis test, $H_3 = 13.16$ , $p = 0.0043$ ), post-hoc tests revealed no significant difference between the sleep-congruent vs. incongruent (two-tailed Mann–Whitney U test, $z = 1.4322$ , $p = 0.2282$ ), wake-congruent vs. incongruent (two-tailed Mann–Whitney U test, $z = 2.1102$ , $p = 0.1044$ ), or sleep-congruent vs. wake-congruent (two-tailed Mann–Whitney U test, $z = 0.00$ , $p = 1.00$ ) groups in the post-interval test session. No significant difference was found for the probe task performance [Probe task (RT), $H_3 = 0.51$ , $p = 0.9172$ ; Probe task (resp), $H_3 = 1.41$ , $p = 0.7024$ ]. All the $p$ -values are not adjusted for multiple comparisons.

**Table S3.**
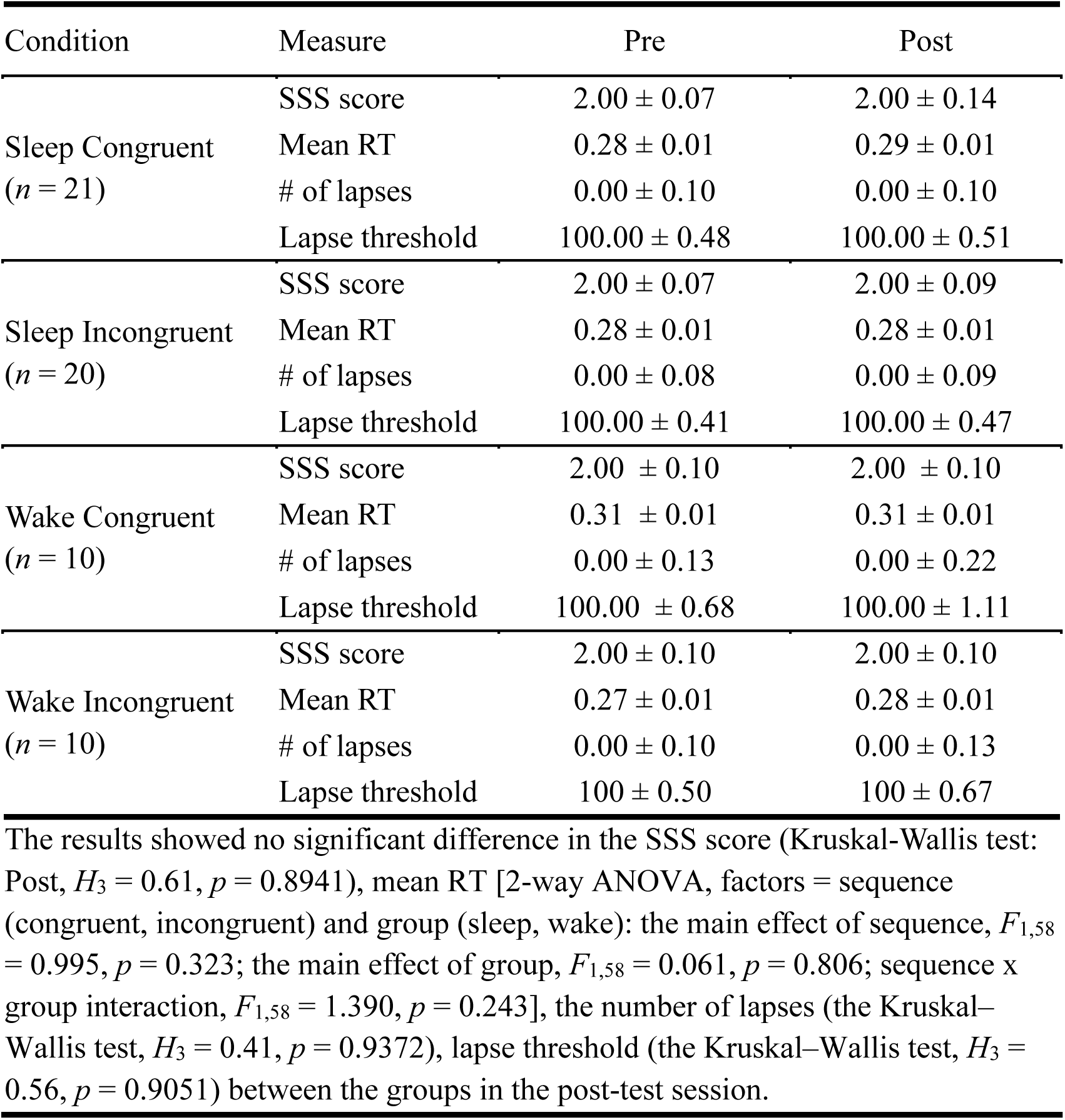
Sleepiness data for experiment 1.

| Condition | Measure | Pre | Post |
| --- | --- | --- | --- |
| Sleep Congruent<br>(*n* = 21) | SSS score | 2.00 ± 0.07 | 2.00 ± 0.14 |
|  | Mean RT | 0.28 ± 0.01 | 0.29 ± 0.01 |
|  | # of lapses | 0.00 ± 0.10 | 0.00 ± 0.10 |
|  | Lapse threshold | 100.00 ± 0.48 | 100.00 ± 0.51 |
| Sleep Incongruent<br>(*n* = 20) | SSS score | 2.00 ± 0.07 | 2.00 ± 0.09 |
|  | Mean RT | 0.28 ± 0.01 | 0.28 ± 0.01 |
|  | # of lapses | 0.00 ± 0.08 | 0.00 ± 0.09 |
|  | Lapse threshold | 100.00 ± 0.41 | 100.00 ± 0.47 |
| Wake Congruent<br>(*n* = 10) | SSS score | 2.00 ± 0.10 | 2.00 ± 0.10 |
|  | Mean RT | 0.31 ± 0.01 | 0.31 ± 0.01 |
|  | # of lapses | 0.00 ± 0.13 | 0.00 ± 0.22 |
|  | Lapse threshold | 100.00 ± 0.68 | 100.00 ± 1.11 |
| Wake Incongruent<br>(*n* = 10) | SSS score | 2.00 ± 0.10 | 2.00 ± 0.10 |
|  | Mean RT | 0.27 ± 0.01 | 0.28 ± 0.01 |
|  | # of lapses | 0.00 ± 0.10 | 0.00 ± 0.13 |
|  | Lapse threshold | 100 ± 0.50 | 100 ± 0.67 |

The results showed no significant difference in the SSS score (Kruskal-Wallis test: Post, $H_3 = 0.61$ , $p = 0.8941$ ), mean RT [2-way ANOVA, factors = sequence (congruent, incongruent) and group (sleep, wake): the main effect of sequence, $F_{1,58} = 0.995$ , $p = 0.323$ ; the main effect of group, $F_{1,58} = 0.061$ , $p = 0.806$ ; sequence x group interaction, $F_{1,58} = 1.390$ , $p = 0.243$ ], the number of lapses (the Kruskal-Wallis test, $H_3 = 0.41$ , $p = 0.9372$ ), lapse threshold (the Kruskal-Wallis test, $H_3 = 0.56$ , $p = 0.9051$ ) between the groups in the post-test session.

**Table S4.** Sleep parameters for experiment 2.

|  | Congruent<br>(*n* = 13) | Incongruent<br>(*n* = 13) | Mann-Whitney U<br>test |  |
| --- | --- | --- | --- | --- |
|  |  |  | *Z* | *P* |
| Sleep efficiency (%) | 66.29 ± 7.42 | 79.06 ± 5.65 | -1.1542 | 0.2484 |
| Wakefulness (%) | 33.35 ± 7.52 | 20.51 ± 5.71 | 1.1803 | 0.2379 |
| NREM 1 (%) | 18.81 ± 3.01 | 20.73 ± 3.04 | -0.4361 | 0.6627 |
| NREM 2(%) | 29.34 ± 5.14 | 41.20 ± 3.34 | -1.5136 | 0.1301 |
| NREM 3 (%) | 15.95 ± 5.46 | 10.98 ± 4.31 | 0.0810 | 0.9355 |
| REM (%) | 2.18 ± 0.84 | 6.15 ± 2.82 | -0.2467 | 0.8052 |
| Time in bed (min) | 86.15 ± 2.66 | 90.00 ± 0.00 | -1.3868 | 0.1655 |

We tested whether the sleep quality between the congruent and incongruent conditions was different, as we directly compared brain activity during sleep between them as discussed in the main text. The statistical results indicated no significant difference in any of the sleep parameters between them. Values are mean ± SEM.

**Table S5.**
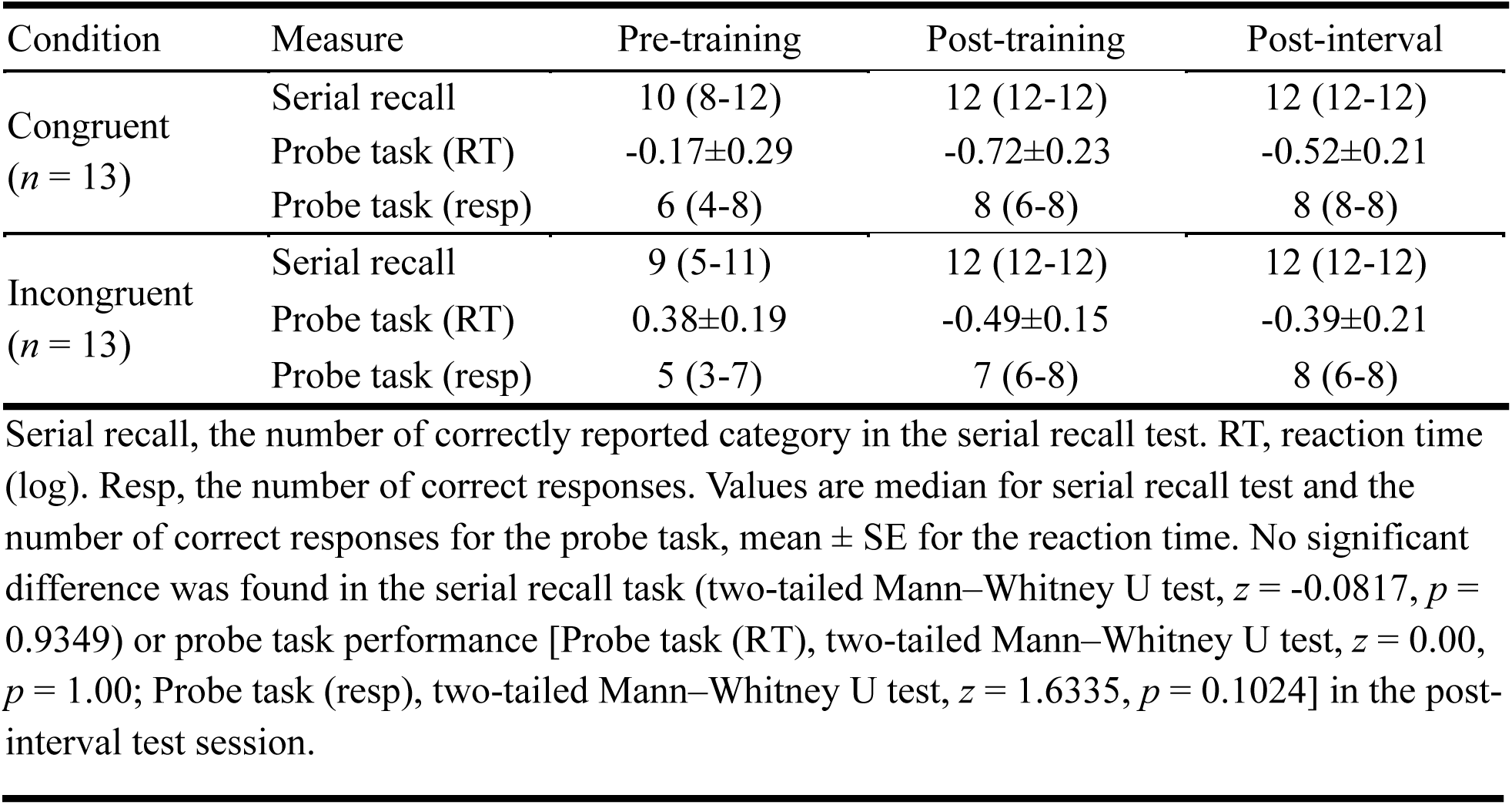
Category task performance for experiment 2.

**Table S6.** MRS data quality for experiment 2.

|  | Congruent<br>(*n* = 12) | Incongruent<br>(*n* = 11) | Mann-Whitney U test |  |
| --- | --- | --- | --- | --- |
|  |  |  | *Z* | *P* |
| Water Linewidth (Hz) | 15.6 ± 0.7 | 17.2 ± 1.2 | -0.4335 | 0.6646 |
| NAA FWHM Linewidth (Hz) | 22.1 ± 1.4 | 21.1 ± 1.8 | 0.4616 | 0.6444 |
| %CRLB for GABA (%) | 6.1 ± 0.6 | 7.2 ± 0.6 | -1.6925 | 0.0905 |
| %CRLB for Glu (%) | 5.8 ± 0.6 | 6.4 ± 0.5 | -0.9540 | 0.3401 |

We compared the MRS quality values between the congruent and incongruent conditions to test whether the MRS data quality was comparable between the groups. The results indicated that there was no systematic bias in the MRS data quality. Thus, the MRS data in these groups were sufficiently reliable for a fair comparison.

**Table S7.**
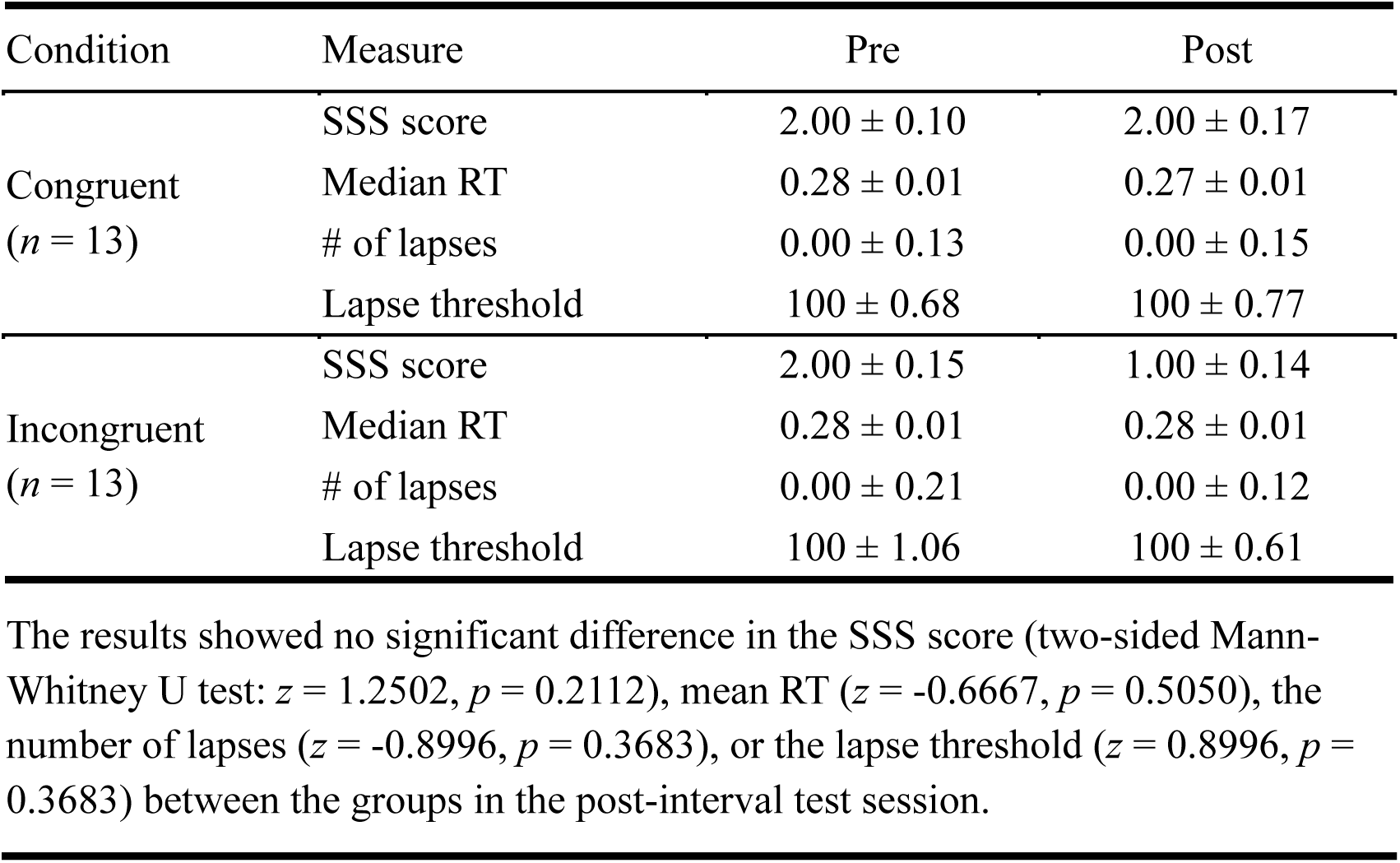
Sleepiness data for experiment 2.

| Condition | Measure | Pre | Post |
| --- | --- | --- | --- |
| Congruent<br>(*n* = 13) | SSS score | 2.00 ± 0.10 | 2.00 ± 0.17 |
|  | Median RT | 0.28 ± 0.01 | 0.27 ± 0.01 |
|  | # of lapses | 0.00 ± 0.13 | 0.00 ± 0.15 |
|  | Lapse threshold | 100 ± 0.68 | 100 ± 0.77 |
| Incongruent<br>(*n* = 13) | SSS score | 2.00 ± 0.15 | 1.00 ± 0.14 |
|  | Median RT | 0.28 ± 0.01 | 0.28 ± 0.01 |
|  | # of lapses | 0.00 ± 0.21 | 0.00 ± 0.12 |
|  | Lapse threshold | 100 ± 1.06 | 100 ± 0.61 |

The results showed no significant difference in the SSS score (two-sided Mann-Whitney U test: $z = 1.2502$ , $p = 0.2112$ ), mean RT ( $z = -0.6667$ , $p = 0.5050$ ), the number of lapses ( $z = -0.8996$ , $p = 0.3683$ ), or the lapse threshold ( $z = 0.8996$ , $p = 0.3683$ ) between the groups in the post-interval test session.

## References

1. W. B. Scoviille, B. Milner, Loss of recent memory after bilateral hippocampal lesions. J. Neurol. Neurosurg. Psychiatry 20, 11–21 (1957).

2. L. R. Squire, Mechanisms of Memory. Science 232, 1612–1619 (1986).

3. E. M. Robertson, Memory leaks: information shared across memory systems. Trends Cogn. Sci. 26, 544–554 (2022).

4. F. Jacobacci, J. L. Armony, A. Yeffal, G. Lerner, E. Amaro, J. Jovicich, J. Doyon, V. Della-Maggiore, Rapid hippocampal plasticity supports motor sequence learning. Proc. Natl. Acad. Sci. 117, 23898–23903 (2020).

5. B. Rasch, J. Born, About Sleep’s Role in Memory. Physiol. Rev. 93, 681–766 (2013).

6. P. Maquet, The Role of Sleep in Learning and Memory. Science 294, 1048–1052 (2001).

7. P. A. Lewis, S. J. Durrant, Overlapping memory replay during sleep builds cognitive schemata. Trends Cogn. Sci. 15, 343–351 (2011).

8. J. M. Ellenbogen, P. T. Hu, J. D. Payne, D. Titone, M. P. Walker, Human relational memory requires time and sleep. Proc. Natl. Acad. Sci. 104, 7723–7728 (2007).

9. N. Dumay, M. G. Gaskell, Sleep-Associated Changes in the Mental Representation of Spoken Words. Psychol. Sci. 18, 35–39 (2007).

10. A. R. Preston, H. Eichenbaum, Interplay of Hippocampus and Prefrontal Cortex in Memory. Curr. Biol. 23, R764–R773 (2013).

11. G. Park, M. S. Kim, Y.-B. Lee, S. Shin, D. Lee, S. J. Kim, Y.-S. Lee, Hippocampal-cortical interactions in the consolidation of social memory. Nat. Commun. 16, 8430 (2025).

12. A. Peyrache, M. Khamassi, K. Benchenane, S. I. Wiener, F. P. Battaglia, Replay of rule-learning related neural patterns in the prefrontal cortex during sleep. Nat. Neurosci. 12, 919–926 (2009).

13. S. Brodt, M. Inostroza, N. Niethard, J. Born, Sleep—A brain-state serving systems memory consolidation. Neuron 111, 1050–1075 (2023).

14. M. Boly, V. Perlbarg, G. Marrelec, M. Schabus, S. Laureys, J. Doyon, M. Pelegrini-Issac, P. Maquet, H. Benali, Hierarchical clustering of brain activity during human nonrapid eye movement sleep. Proc. Natl. Acad. Sci. 109, 5856–5861 (2012).

15. M. Massimini, F. Ferrarelli, R. Huber, S. K. Esser, H. Singh, G. Tononi, Breakdown of Cortical Effective Connectivity During Sleep. Science 309, 2228–2232 (2005).

16. B. P. Staresina, Coupled sleep rhythms for memory consolidation. Trends Cogn. Sci. 28, 339–351 (2024).

17. P. Simor, G. van der Wijk, L. Nobili, P. Peigneux, The microstructure of REM sleep: Why phasic and tonic? Sleep Med. Rev. 52, 101305 (2020).

18. N. Mosha, E. M. Robertson, Unstable Memories Create a High-Level Representation that Enables Learning Transfer. Curr. Biol. 26, 100–105 (2016).

19. K. Shibata, Y. Sasaki, J. W. Bang, E. G. Walsh, M. G. Machizawa, M. Tamaki, L.-H. Chang, T. Watanabe, Overlearning hyperstabilizes a skill by rapidly making neurochemical processing inhibitory-dominant. Nat. Neurosci. 20, 470–475 (2017).

20. Y. Dudai, The Neurobiology of Consolidations, Or, How Stable is the Engram? Annu. Rev. Psychol. 55, 51–86 (2004).

21. G. Tononi, C. Cirelli, Sleep and the price of plasticity: from synaptic and cellular homeostasis to memory consolidation and integration. Neuron 81, 12–34 (2014).

22. J. G. Klinzing, N. Niethard, J. Born, Mechanisms of systems memory consolidation during sleep. Nat. Neurosci. 22, 1598–1610 (2019).

23. M. Tamaki, Z. Wang, T. Barnes-Diana, D. Guo, A. V. Berard, E. Walsh, T. Watanabe, Y. Sasaki, Complementary contributions of non-REM and REM sleep to visual learning. Nat. Neurosci. 23, 1150–1156 (2020).

24. M. Tamaki, A. V. Berard, T. Barnes-Diana, J. Siegel, T. Watanabe, Y. Sasaki, Reward does not facilitate visual perceptual learning until sleep occurs. Proc. Natl. Acad. Sci. 117, 959–968 (2020).

25. D. G. Almeida-Filho, C. M. Queiroz, S. Ribeiro, Memory corticalization triggered by REM sleep: mechanisms of cellular and systems consolidation. Cell. Mol. Life Sci. 75, 3715–3740 (2018).

26. D. G. de Almeida-Filho, B. D. V. Koike, F. Billwiller, K. S. Farias, I. R. P. de Sales, P.-H. Luppi, S. Ribeiro, C. M. Queiroz, Hippocampus-retrosplenial cortex interaction is increased during phasic REM and contributes to memory consolidation. Sci. Rep. 11, 13078 (2021).

27. R. Boyce, S. D. Glasgow, S. Williams, A. Adamantidis, Causal evidence for the role of REM sleep theta rhythm in contextual memory consolidation. Science 352, 812–816 (2016).

28. B. Rasch, C. Buchel, S. Gais, J. Born, Odor Cues During Slow-Wave Sleep Prompt Declarative Memory Consolidation. Science 315, 1426–1429 (2007).

29. J. D. Rudoy, J. L. Voss, C. E. Westerberg, K. A. Paller, Strengthening Individual Memories by Reactivating Them During Sleep. Science 326, 1079–1079 (2009).

30. T. K. Hensch, Critical period plasticity in local cortical circuits. Nat. Rev. Neurosci. 6, 877–888 (2005).

31. B. M. Hooks, C. Chen, Critical Periods in the Visual System: Changing Views for a Model of Experience-Dependent Plasticity. Neuron 56, 312–326 (2007).

32. J. W. Bang, K. Shibata, S. M. Frank, E. G. Walsh, M. W. Greenlee, T. Watanabe, Y. Sasaki, Consolidation and reconsolidation share behavioural and neurochemical mechanisms. *Nat*. Hum. Behav. 2, 507–513 (2018).

33. T. Yamada, T. Watanabe, Y. Sasaki, Plasticity–stability dynamics during post-training processing of learning. Trends Cogn. Sci. 28, 72–83 (2023).

34. J. A. Stanley, N. Raz, Functional Magnetic Resonance Spectroscopy: The “New” MRS for Cognitive Neuroscience and Psychiatry Research. Front. Psychiatry 9, 76 (2018).

35. M. Tamaki, T. Watanabe, Y. Sasaki, Coregistration of magnetic resonance spectroscopy and polysomnography for sleep analysis in human subjects. STAR Protoc. 2, 100974 (2021).

36. M. Mescher, H. Merkle, J. Kirsch, M. Garwood, R. Gruetter, Simultaneous in vivo spectral editing and water suppression. NMR Biomed. 11, 7 (1998).

37. D.-J. Dijk, EEG slow waves and sleep spindles: windows on the sleeping brain. Behav. Brain Res. 69, 109–116 (1995).

38. F. Amzica, M. Steriade, Electrophysiological correlates of sleep delta waves. Electroencephalogr. Clin. Neurophysiol. 107, 69–83 (1998).

39. A. Karni, G. Meyer, P. Jezzard, M. M. Adams, R. Turner, L. G. Ungerleider, Functional MRI evidence for adult motor cortex plasticity during motor skill learning. Nature 377, 155–158 (1995).

40. A. Karni, G. Meyer, C. Rey-Hipolito, P. Jezzard, M. M. Adams, R. Turner, L. G. Ungerleider, The acquisition of skilled motor performance: Fast and slow experience-driven changes in primary motor cortex. Proc. Natl. Acad. Sci. 95, 861–868 (1998).

41. V. Sterpenich, C. Schmidt, G. Albouy, L. Matarazzo, A. Vanhaudenhuyse, P. Boveroux, C. Degueldre, Y. Leclercq, E. Balteau, F. Collette, A. Luxen, C. Phillips, P. Maquet, Memory Reactivation during Rapid Eye Movement Sleep Promotes Its Generalization and Integration in Cortical Stores. Sleep 37, 1061–1075 (2014).

42. S. I. R. Pereira, L. Santamaria, R. Andrews, E. Schmidt, M. C. W. Van Rossum, P. Lewis, Rule Abstraction Is Facilitated by Auditory Cuing in REM Sleep. J. Neurosci. 43, 3838–3848 (2023).

43. C.-F. V. Latchoumane, H.-V. V. Ngo, J. Born, H.-S. Shin, Thalamic Spindles Promote Memory Formation during Sleep through Triple Phase-Locking of Cortical, Thalamic, and Hippocampal Rhythms. Neuron 95, 424–435.e6 (2017).

44. S. A. Cairney, A. Á. V. Guttesen, N. El Marj, B. P. Staresina, Memory Consolidation Is Linked to Spindle-Mediated Information Processing during Sleep. Curr. Biol. 28, 948–954.e4 (2018).

45. A. A. Perrault, A. Khani, C. Quairiaux, K. Kompotis, P. Franken, M. Muhlethaler, S. Schwartz, L. Bayer, Whole-Night Continuous Rocking Entrains Spontaneous Neural Oscillations with Benefits for Sleep and Memory. Curr. Biol. 29, 402–411.e3 (2019).

46. P. Peigneux, S. Laureys, S. Fuchs, A. Destrebecqz, F. Collette, X. Delbeuck, C. Phillips, J. Aerts, G. Del Fiore, C. Degueldre, A. Luxen, A. Cleeremans, P. Maquet, Learned material content and acquisition level modulate cerebral reactivation during posttraining rapid-eye-movements sleep. NeuroImage 20, 125–134 (2003).

47. J. Kim, A. Joshi, L. Frank, K. Ganguly, Cortical–hippocampal coupling during manifold exploration in motor cortex. Nature 613, 103–110 (2023).

48. M. Nishida, M. P. Walker, Daytime Naps, Motor Memory Consolidation and Regionally Specific Sleep Spindles. PLoS ONE 2, e341 (2007).

49. M. Tamaki, T.-R. Huang, Y. Yotsumoto, M. Hamalainen, F.-H. Lin, J. E. Nanez, T. Watanabe, Y. Sasaki, Enhanced Spontaneous Oscillations in the Supplementary Motor Area Are Associated with Sleep-Dependent Offline Learning of Finger-Tapping Motor-Sequence Task. J. Neurosci. 33, 13894–13902 (2013).

50. D. S. Manoach, K. N. Thakkar, E. Stroynowski, A. Ely, S. K. McKinley, E. Wamsley, I. Djonlagic, M. G. Vangel, D. C. Goff, R. Stickgold, Reduced overnight consolidation of procedural learning in chronic medicated schizophrenia is related to specific sleep stages. J. Psychiatr. Res. 44, 112–120 (2010).

51. B. Crosson, Activity in the Paracingulate and Cingulate Sulci during Word Generation: An fMRI Study of Functional Anatomy. Cereb. Cortex 9, 307–316 (1999).

52. N. Picard, P. L. Strick, Imaging the premotor areas. Curr. Opin. Neurobiol. 11, 663–672 (2001).

53. A. Yokoi, J. Diedrichsen, Neural Organization of Hierarchical Motor Sequence Representations in the Human Neocortex. Neuron 103, 1178–1190.e7 (2019).

54. S. A. Bunge, I. Kahn, J. D. Wallis, E. K. Miller, A. D. Wagner, Neural Circuits Subserving the Retrieval and Maintenance of Abstract Rules. J. Neurophysiol. 90, 3419–3428 (2003).

55. H. Eichenbaum, Prefrontal–hippocampal interactions in episodic memory. Nat. Rev. Neurosci. 18, 547–558 (2017).

56. C. R. Bowman, D. Zeithamova, Abstract Memory Representations in the Ventromedial Prefrontal Cortex and Hippocampus Support Concept Generalization. J. Neurosci. 38, 2605–2614 (2018).

57. R. Wehrle, C. Kaufmann, T. C. Wetter, F. Holsboer, D. P. Auer, T. Pollmächer, M. Czisch, Functional microstates within human REM sleep: first evidence from fMRI of a thalamocortical network specific for phasic REM periods: Thalamocortical network in phasic REM sleep. Eur. J. Neurosci. 25, 863–871 (2007).

58. M. Tamaki, Y. Sasaki, Surveillance during REM sleep for the first-night effect. Front. Neurosci. 13, 1161 (2019).

59. N. H. Van Den Berg, A. Gibbings, D. Baena, A. Pozzobon, J. Al-Kuwatli, L. B. Ray, S. M. Fogel, Eye movements during phasic versus tonic rapid eye movement sleep are biomarkers of dissociable electroencephalogram processes for the consolidation of novel problem-solving skills. SLEEP 46, zsad151 (2023).

60. A. A. Borbely, A Two Process Model of Sleep Regulation. Hum. Neurobiol. 1, 195–204 (1982).

61. P. Franken, D.-J. Dijk, Sleep and circadian rhythmicity as entangled processes serving homeostasis. Nat. Rev. Neurosci. 25, 43–59 (2024).

62. R. Scheeringa, P. Fries, K.-M. Petersson, R. Oostenveld, I. Grothe, D. G. Norris, P. Hagoort, M. C. M. Bastiaansen, Neuronal Dynamics Underlying High- and Low-Frequency EEG Oscillations Contribute Independently to the Human BOLD Signal. Neuron 69, 572–583 (2011).

63. M. Uji, R. Wilson, S. T. Francis, K. J. Mullinger, S. D. Mayhew, Exploring the advantages of multiband fMRI with simultaneous EEG to investigate coupling between gamma frequency neural activity and the BOLD response in humans. Hum. Brain Mapp. 39, 1673–1687 (2018).

64. E. A. Woodcock, C. Anand, D. Khatib, V. A. Diwadkar, J. A. Stanley, Working Memory Modulates Glutamate Levels in the Dorsolateral Prefrontal Cortex during 1H fMRS. Front. Psychiatry 9, 66 (2018).

65. V. Bezalel, R. Paz, A. Tal, Inhibitory and excitatory mechanisms in the human cingulate-cortex support reinforcement learning: A functional Proton Magnetic Resonance Spectroscopy study. NeuroImage 184, 25–35 (2019).

66. N. Niethard, H.-V. V. Ngo, I. Ehrlich, J. Born, Cortical circuit activity underlying sleep slow oscillations and spindles. Proc. Natl. Acad. Sci. 115 (2018).

67. N. Niethard, S. Brodt, J. Born, Cell-Type-Specific Dynamics of Calcium Activity in Cortical Circuits over the Course of Slow-Wave Sleep and Rapid Eye Movement Sleep. J. Neurosci. 41, 4212–4222 (2021).

68. R. Huber, M. Felice Ghilardi, M. Massimini, G. Tononi, Local sleep and learning. Nature 430, 78–81 (2004).

69. G. Tononi, C. Cirelli, Sleep and synaptic homeostasis: a hypothesis. Brain Res. Bull. 62, 143–150 (2003).

70. H. Norimoto, K. Makino, M. Gao, Y. Shikano, K. Okamoto, T. Ishikawa, T. Sasaki, H. Hioki, S. Fujisawa, Y. Ikegaya, Hippocampal ripples down-regulate synapses. Science 359, 1524–1527 (2018).

71. A. Giuditta, M. V. Ambrosini, P. Montagnese, P. Mandile, M. Cotugno, G. G. Zucconi, S. Vescia, The sequential hypothesis of the function of sleep. Behav. Brain Res. 69, 157–166 (1995).

72. A. Giuditta, Sleep memory processing: the sequential hypothesis. Front. Syst. Neurosci. 8 (2014).

73. M. V. Ambrosini, M. Langella, U. A. Gironi Carnevale, A. Giuditta, The sequential hypothesis of sleep function. III. The structure of postacquisition sleep in learning and nonlearning rats. Physiol. Behav. 51, 217–226 (1992).

74. Y. Dudai, The Restless Engram: Consolidations Never End. Annu. Rev. Neurosci. 35, 227–247 (2012).

75. L. Bastian, A. Samanta, D. Ribeiro De Paula, F. D. Weber, R. Schoenfeld, M. Dresler, L. Genzel, Spindle–slow oscillation coupling correlates with memory performance and connectivity changes in a hippocampal network after sleep. Hum. Brain Mapp. 43, 3923–3943 (2022).

76. C. Kjaerby, M. Andersen, N. Hauglund, V. Untiet, C. Dall, B. Sigurdsson, F. Ding, J. Feng, Y. Li, P. Weikop, H. Hirase, M. Nedergaard, Memory-enhancing properties of sleep depend on the oscillatory amplitude of norepinephrine. Nat. Neurosci. 25, 1059–1070 (2022).

77. T. Xia, D. Chen, S. Zeng, Z. Yao, J. Liu, S. Qin, K. A. Paller, S. G. Torres Platas, J. W. Antony, X. Hu, Aversive memories can be weakened during human sleep via the reactivation of positive interfering memories. Proc. Natl. Acad. Sci. 121, e2400678121 (2024).

78. D. Oudiette, K. A. Paller, Upgrading the sleeping brain with targeted memory reactivation. Trends Cogn. Sci. 17, 142–149 (2013).

79. B. A. Mander, J. R. Winer, M. P. Walker, Sleep and Human Aging. Neuron 94, 19–36 (2017).

80. Y. Takado, H. Takuwa, K. Sampei, T. Urushihata, M. Takahashi, M. Shimojo, S. Uchida, N. Nitta, S. Shibata, K. Nagashima, Y. Ochi, M. Ono, J. Maeda, Y. Tomita, N. Sahara, J. Near, I. Aoki, K. Shibata, M. Higuchi, MRS-measured glutamate versus GABA reflects excitatory versus inhibitory neural activities in awake mice. J. Cereb. Blood Flow Metab. 42, 197–212 (2022).

81. D. Pasanta, J. L. He, T. Ford, G. Oeltzschner, D. J. Lythgoe, N. A. Puts, Functional MRS studies of GABA and glutamate/Glx – A systematic review and meta-analysis. Neurosci. Biobehav. Rev. 144, 104940 (2023).

82. R. F. Helfrich, J. D. Lendner, R. T. Knight, Aperiodic sleep networks promote memory consolidation. Trends Cogn. Sci. 25, 648–659 (2021).

83. J. D. Lendner, N. Niethard, B. A. Mander, F. J. van Schalkwijk, S. Schuh-Hofer, H. Schmidt, R. T. Knight, J. Born, M. P. Walker, J. J. Lin, R. F. Helfrich, Human REM sleep recalibrates neural activity in support of memory formation. Sci. Adv. 9, eadj1895 (2023).

84. A. Cochrane, L. Rosedahl, M. Tamaki, T. Watanabe, Y. Sasaki, Concurrent multimodal imaging demonstrates that EEG-based excitation/inhibition balance reflects glutamate and GABA concentrations. doi: 10.1523/JNEUROSCI.1394-25.2026 (2026).

85. F. Faul, E. Erdfelder, A.-G. Lang, A. Buchner, G*Power 3: A flexible statistical power analysis program for the social, behavioral, and biomedical sciences. Behav. Res. Methods 39, 175–191 (2007).

86. M. Tamaki, Z. Wang, T. Watanabe, Y. Sasaki, Trained-feature–specific offline learning by sleep in an orientation detection task. J. Vis. 19, 12 (2019).

87. M. Tamaki, J. W. Bang, T. Watanabe, Y. Sasaki, The first-night effect suppresses the strength of slow-wave activity originating in the visual areas during sleep. Vision Res. 99, 154–161 (2014).

88. M. Uji, X. Li, A. Saotome, R. Katsumata, R. A. Waggoner, C. Suzuki, K. Ueno, S. Aritake, M. Tamaki, Human deep sleep facilitates cerebrospinal fluid dynamics linked to spontaneous brain oscillations and neural events. Proc. Natl. Acad. Sci. 122, e2509626122 (2025).

89. D. Tomasi, N. D. Volkow, Associations between handedness and brain functional connectivity patterns in children. Nat. Commun. 15, 2355 (2024).

90. H. W. Agnew, W. W. Webb, R. L. Williams, Sleep patterns in late middle age males: An EEG study. Electroencephalogr. Clin. Neurophysiol. 23, 168–171 (1967).

91. M. Tamaki, H. Nittono, M. Hayashi, T. Hori, Examination of the First-Night Effect during the Sleep-Onset Period. Sleep 28, 195–202 (2005).

92. M. Tamaki, H. Nittono, T. Hori, The first-night effect occurs at the sleep-onset period regardless of the temporal anxiety level in healthy students. Sleep Biol. Rhythms 3, 92–94 (2005).

93. M. Tamaki, J. W. Bang, T. Watanabe, Y. Sasaki, Night Watch in One Brain Hemisphere during Sleep Associated with the First-Night Effect in Humans. Curr. Biol. 26, 1190–1194 (2016).

94. J. W. Bang, O. Khalilzadeh, M. Hämäläinen, T. Watanabe, Y. Sasaki, Location specific sleep spindle activity in the early visual areas and perceptual learning. Vision Res. 99, 162–171 (2014).

95. M. P. Walker, Sleep and the Time Course of Motor Skill Learning. Learn. Mem. 10, 275–284 (2003).

96. K. Kuriyama, Sleep-dependent learning and motor-skill complexity. Learn. Mem. 11, 705–713 (2004).

97. E. Hoddes, V. Zarcone, H. Smythe, R. Phillips, W. C. Dement, Quantification of Sleepiness: A New Approach. Psychophysiology 10, 431–436 (1973).

98. D. F. Dinges, J. W. Powell, Microcomputer analyses of performance on a portable, simple visual RT task during sustained operations. Behav. Res. Methods Instrum. Comput. 17, 652–655 (1985).

99. J. Lim, D. F. Dinges, Sleep Deprivation and Vigilant Attention. Ann. N. Y. Acad. Sci. 1129, 305–322 (2008).

100. J. Frahm, K.-D. Merboldt, W. Hänicke, Localized proton spectroscopy using stimulated echoes. J. Magn. Reson. 1969 72, 502–508 (1987).

101. A. Haase, J. Frahm, W. Hanicke, D. Matthaei, ^1^ H NMR chemical shift selective (CHESS) imaging. Phys. Med. Biol. 30, 341–344 (1985).

102. C. Gasparovic, T. Song, D. Devier, H. J. Bockholt, A. Caprihan, P. G. Mullins, S. Posse, R. E. Jung, L. A. Morrison, Use of tissue water as a concentration reference for proton spectroscopic imaging. Magn. Reson. Med. 55, 1219–1226 (2006).

103. G. Oeltzschner, A. Schnitzler, F. Wickrath, H. J. Zöllner, H.-J. Wittsack, Use of quantitative brain water imaging as concentration reference for J-edited MR spectroscopy of GABA. Magn. Reson. Imaging 34, 1057–1063 (2016).

104. W. T. Clarke, C. J. Stagg, S. Jbabdi, FSL-MRS: An end-to-end spectroscopy analysis package. Magn. Reson. Med. 85, 2950–2964 (2021).

105. P. J. Allen, O. Josephs, R. Turner, A method for removing imaging artifact from continuous EEG recorded during functional MRI. NeuroImage 12, 230–239 (2000).

106. R. Wilson, K. J. Mullinger, S. T. Francis, S. D. Mayhew, The relationship between negative BOLD responses and ERS and ERD of alpha/beta oscillations in visual and motor cortex. NeuroImage 199, 635–650 (2019).

107. T. Warbrick, Simultaneous EEG-fMRI: What Have We Learned and What Does the Future Hold? Sensors 22, 2262 (2022).

108. M. Bullock, G. D. Jackson, D. F. Abbott, Artifact Reduction in Simultaneous EEG-fMRI: A Systematic Review of Methods and Contemporary Usage. Front. Neurol. 12 (2021).

109. A. Rechtschaffen, A. Kales, *A Manual of Standardized Terminology, Techniques and Scoring System for Sleep Stages of Human Subjects*. (Public Health Service, US Government Printing Office, Washington DC., 1968).

110. R. Berry, R. Brooks, C. Gamaldo, S. Harding, R. Lloyd, S. Quan, M. Troester, B. Vaughn, AASM scoring manual updates for 2017 (Version 2.4). J Clin Sleep Med 13, 665–666 (2017).

111. M. J. Brookes, J. Vrba, K. J. Mullinger, G. B. Geirsdóttir, W. X. Yan, C. M. Stevenson, R. Bowtell, P. G. Morris, Source localisation in concurrent EEG/fMRI: Applications at 7T. NeuroImage 45, 440–452 (2009).

112. M. Uji, N. Cross, F. B. Pomares, A. A. Perrault, A. Jegou, A. Nguyen, U. Aydin, J.-M. Lina, T. T. Dang-Vu, C. Grova, Data-driven beamforming technique to attenuate ballistocardiogram artefacts in electroencephalography-functional magnetic resonance imaging without detecting cardiac pulses in electrocardiography recordings. Hum. Brain Mapp. 42, 3993–4021 (2021).

113. B. D. Van Veen, W. van Drongelen, M. Yuchtman, A. Suzuki, Localization of brain electrical activity via linearly constrained minimum variance spatial filtering. IEEE Trans. Biomed. Eng. 44, 867–880 (1997).

114. F. Tadel, S. Baillet, J. C. Mosher, D. Pantazis, R. M. Leahy, Brainstorm: a user-friendly application for MEG/EEG analysis. Comput. Intell. Neurosci. 2011, 879716 (2011).

115. V. Fonov, A. Evans, R. McKinstry, C. Almli, D. Collins, Unbiased nonlinear average age-appropriate brain templates from birth to adulthood. NeuroImage 47, S102 (2009).

116. V. Fonov, A. C. Evans, K. Botteron, C. R. Almli, R. C. McKinstry, D. L. Collins, Unbiased average age-appropriate atlases for pediatric studies. NeuroImage 54, 313–327 (2011).

117. J. Kybic, M. Clerc, O. Faugeras, R. Keriven, T. Papadopoulo, Fast multipole acceleration of the MEG/EEG boundary element method. Phys. Med. Biol. 50, 4695 (2005).

118. J. Zhang, G. Miao, A linear hybrid model of MSE and BEM for floating structures in coastal zones1. J. Hydrodyn. Ser B 18, 649–658 (2006).

119. R. A. Poldrack, J. Clark, E. J. Paré-Blagoev, D. Shohamy, J. Creso Moyano, C. Myers, M. A. Gluck, Interactive memory systems in the human brain. Nature 414, 546–550 (2001).

120. E. T. Rolls, C.-C. Huang, C.-P. Lin, J. Feng, M. Joliot, Automated anatomical labelling atlas 3. NeuroImage 206, 116189 (2020).

121. M. Massimini, R. Huber, F. Ferrarelli, S. Hill, G. Tononi, The Sleep Slow Oscillation as a Traveling Wave. J. Neurosci. 24, 6862–6870 (2004).

122. M. Mölle, T. O. Bergmann, L. Marshall, J. Born, Fast and Slow Spindles during the Sleep Slow Oscillation: Disparate Coalescence and Engagement in Memory Processing. Sleep 34, 1411–1421 (2011).

123. C. L. Scrivener, A. T. Reader, Variability of EEG electrode positions and their underlying brain regions: visualizing gel artifacts from a simultaneous EEG-fMRI dataset. Brain Behav. 12, e2476 (2022).

124. T. Schreiner, E. Kaufmann, S. Noachtar, J.-H. Mehrkens, T. Staudigl, The human thalamus orchestrates neocortical oscillations during NREM sleep. Nat. Commun. 13, 5231 (2022).

125. D. Baena, E. Gabitov, L. B. Ray, J. Doyon, S. M. Fogel, Motor learning promotes regionally-specific spindle-slow wave coupled cerebral memory reactivation. *Commun*. Biol. 7, 1492 (2024).

126. R. Agarwal, T. Takeuchi, S. Laroche, J. Gotman, Detection of rapid-eye movements in sleep studies. IEEE Trans. Biomed. Eng. 52, 1390–1396 (2005).

127. R. Oostenveld, P. Fries, E. Maris, J.-M. Schoffelen, FieldTrip: Open source software for advanced analysis of MEG, EEG, and invasive electrophysiological data. Comput. Intell. Neurosci. 2011, 156869 (2011).

128. W. T. Clarke, C. J. Stagg, S. Jbabdi, FSL-MRS: An end-to-end spectroscopy analysis package. Magn. Reson. Med. 85, 2950–2964 (2021).

129. S. W. Provencher, Automatic quantitation of localizedin vivo1H spectra with LCModel. NMR Biomed. 14, 260–264 (2001).

130. J. Near, A. D. Harris, C. Juchem, R. Kreis, M. Marjańska, G. Öz, J. Slotboom, M. Wilson, C. Gasparovic, Preprocessing, analysis and quantification in single-voxel magnetic resonance spectroscopy: experts’ consensus recommendations. NMR Biomed. 34, e4257 (2021).

131. N. Bloembergen, E. M. Purcell, R. V. Pound, Relaxation Effects in Nuclear Magnetic Resonance Absorption. Phys. Rev. 73, 679–712 (1948).

132. R. Kreis, Issues of spectral quality in clinical1H-magnetic resonance spectroscopy and a gallery of artifacts. NMR Biomed. 17, 361–381 (2004).

133. Y. Benjamini, Y. Hochberg, Controlling the False Discovery Rate: A Practical and Powerful Approach to Multiple Testing. J. R. Stat. Soc. Ser. B Methodol. 57, 289–300 (1995).

